# Estimating recent migration and population size surfaces

**DOI:** 10.1101/365536

**Authors:** Hussein Al-Asadi, Desislava Petkova, Matthew Stephens, John Novembre

**Author notes:** to whom correspondence shall be addressed. July 9, 2018.

## Abstract

In many species a fundamental feature of genetic diversity is that genetic similarity decays with geographic distance; however, this relationship is often complex, and may vary across space and time. Methods to uncover and visualize such relationships have widespread use for analyses in molecular ecology, conservation genetics, evolutionary genetics, and human genetics. While several frameworks exist, a promising approach is to infer maps of how migration rates vary across geographic space. Such maps could, in principle, be estimated across time to reveal the full complexity of population histories. Here, we take a step in this direction: we present a method to infer separate maps of population sizes and migration rates for different time periods from a matrix of genetic similarity between every pair of individuals. Specifically, genetic similarity is measured by counting the number of long segments of haplotype sharing (also known as identity-by-descent tracts). By varying the length of these segments we obtain parameter estimates for qualitatively different time periods. Using simulations, we show that the method can reveal time-varying migration rates and population sizes, including changes that are not detectable when ignoring haplotypic structure. We apply the method to a dataset of contemporary European individuals (POPRES), and provide an integrated analysis of recent population structure and growth over the last *~*3,000 years in Europe. Software implementing the methods is available at https://github.com/halasadi/MAPS.

## 1 Introduction

Populations exist on a physical landscape and often have limited dispersal. As a result, most genetic data exhibit a pattern of isolation by distance (Wright, 1943), which is simply to say that populations closer to each other geographically are more similar genetically. Furthermore, the degree of isolation by distance can vary across space and time (Manel et al., 2003). For instance, in a mountainous area of a terrestrial species’ range, a pair of individuals may be more divergent from each other than a pair of individuals separated by the same distance in a flat and open area of the habitat. Additionally, the degree of isolation by distance can change over time – for example, if dispersal patterns are changing over time. Such spatial and temporal heterogeneity is an important aspect of population biology, and understanding it is crucial to solving problems in ecology (Turner et al., 2001), conservation genetics (Segelbacher et al., 2010), evolution (Rousset, 2004), and human genetics (Rosenberg et al., 2005).

Several methods have been developed to reveal spatial heterogeneity in patterns of isolation by distance (Womble, 1951; Barbujani et al., 1989; Guillot et al., 2005, 2009; Caye et al., 2016; Petkova et al., 2016; Bradburd et al., 2016, 2017). Some methods are based on explicitly modeling the spatial structure in the data (Guillot et al., 2005, 2009; Petkova et al., 2016; Bradburd et al., 2016, 2017); others take non-parametric approaches (e.g. Womble, 1951; Barbujani et al., 1989); while other methods ignore the spatial configuration of the samples and rely on researchers to make a *post hoc* geographic interpretation of the results (e.g. Pritchard et al., 2000; Patterson et al., 2006). However, none of these methods can be flexibly applied to address temporal heterogeneity in isolation by distance patterns, and new methods are needed.

One source of information for inferring changes in demography across time is the density of mutations observed in pairwise sequence comparisons (Li and Durbin, 2011; Schraiber and Akey, 2015). For example, when individuals are similar along a long segment of their chromosomes, it suggests that these segments share a recent common ancestor (Palamara et al., 2012). These segments are often called “identity-by-descent” tracts, although here we prefer the term “long pairwise shared coalescence” (lPSC) segments (as identity by descent traditionally required a definition of a founder generation, which is not clear in most data applications). A key feature of these segments is that filtering them by length provides a means to interrogate different periods of population history. The longest segments reflect the most recent population history, whereas shorter segments reflect longer periods of time. Recent analyses using lPSC segments suggest that they can reveal fine-scale spatial and temporal patterns of population structure that are not evident with genotype-based methods such as principal components analysis (Ralph and Coop, 2013; Lawson et al., 2012; Leslie et al., 2015).

Here we develop a new method to infer spatial and temporal heterogeneity in population sizes and migration rates. The method takes as input geographic coordinates for a set of individuals sampled across a spatial landscape, and a matrix of their genetic similarities as measured by sharing of lPSC segments. It then infers two maps, one representing dispersal rates across the landscape, and another representing population density. Crucially, building these maps using different lengths of lPSC segments can help reveal changes in dispersal rates and population sizes over time.

Our method is based on a stepping-stone model where randomly-mating subpopulations are connected to neighboring subpopulations in a grid. Such models are parameterized by a vector of population sizes (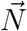) and a sparse migration rate matrix (**M**). Stepping-stone models with a large number of demes can approximate spatially continuous population models (Barton et al., 2002; Baharian et al., 2016), and this can be exploited to produce maps of approximate dispersal rates and population density across continuous space.

Our method builds upon a method developed for estimating effective migration surfaces (EEMS) (Petkova et al., 2016). While EEMS infers local rates of effective migration relative to a global average, here we can explicitly infer absolute parameter values by leveraging lPSC segments and modeling the recombination process [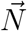 and **M** values in the stepping-stone model, and effective spatial density function *D_e_*(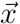) and dispersal rate function *σ*(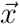) in the continuous limit]. We call this method MAPS, for inferring Migration And Population-size Surfaces.

We test MAPS on coalescent simulations and apply it to a European subset of 2,224 individuals from the POPRES data (Nelson et al., 2008). In simulations, we show that MAPS can infer both time-resolved migration barriers and population sizes across the habitat. In empirical data, we infer dispersal rates *σ*(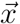) and population densities *D_e_*(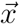) across different time periods in Europe.

## 2 Results

### 2.1 Outline of the MAPS method

MAPS estimates demography using the number of Pairwise Shared Coalescence (PSC) segments of different lengths shared between individuals. We define a PSC segment between (haploid) individuals to be a genomic segment with a single coalescent time across its length (Figure 1A). Long PSC (lPSC) segments tend to have a recent coalescent time, and so manifest themselves in genotype data as unusually long regions of high pairwise similarity, which can be detected by various software packages (Gusev et al., 2009; Browning and Browning, 2011, 2013; Chiang et al., 2016). Because lPSC segments typically reflect recent coalescent events, counts of lPSC segments are especially informative for recent population structure (Ringbauer et al., 2017; Palamara et al., 2012; Baharian et al., 2016). And partitioning lPSC segments into different lengths bins (e.g. 2-8cM, *≥*8cM) can help focus inference on different (recent) temporal scales.

**Figure 1:**
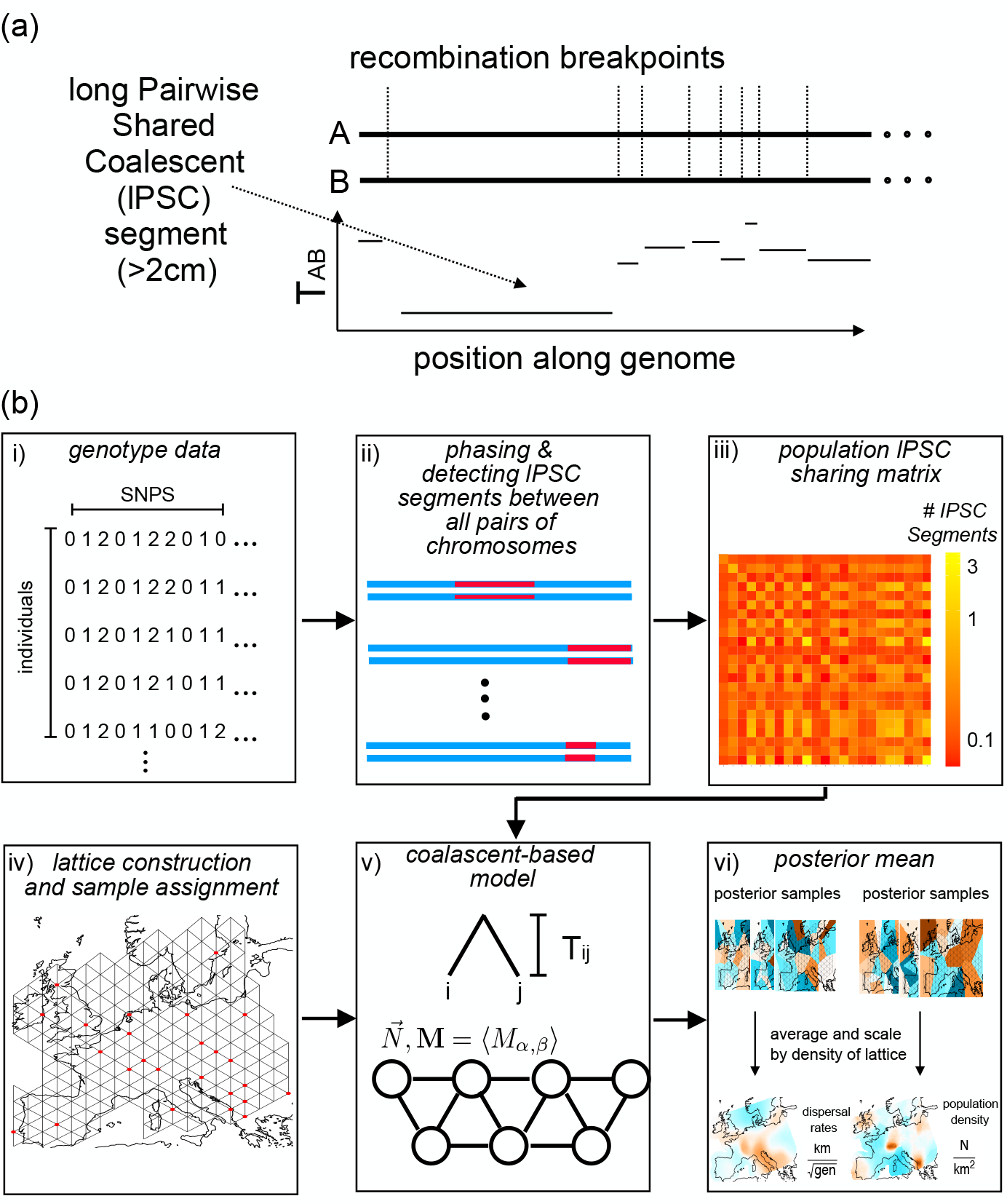
Schematic overview of MAPS. (a) Coalescent times between a pair of hapolo-types (A and B) will vary across the genome in discrete segments bordered by recombination breakpoints. On average, longer segments represent shorter pairwise coalescent times (*T_AB_*) (b) Flow diagram of MAPS. i) We start with a matrix of called genotypes; ii) lPSC segments between all pairs of chromosomes across the genome are identified from the data using external methods (such as BEAGLE, Browning and Browning (2011)); iii) lPSC segments between pairs of individuals are aggregated at the levels of pairs of populations; iv) A grid is constructed and individuals are assigned to the most nearby node; v) The probability of the PSC sharing matrix can be computed under a stepping-stone model where each node represents a population and each edge represents symmetric migration; vi) We use an MCMC scheme to sample from the posterior distribution of migration rates and population sizes. The final MAPS output is the mean over these posterior samples, and the averaged rates can be transformed to units of dispersal rate and population density. The diagram does not show a bootstrapping step used to estimate likelihood weights to account for correlations between lPSC segments, see Equation (6) in Methods.

The MAPS model involves two components: i) a likelihood function, which relates the observed data (genetic similarities, as measured by sharing of lPSC segments) to the underlying demographic parameters (migration rates and population sizes); and ii) a prior distribution on the demographic parameters, which captures the idea that nearby locations will often have similar demographic parameters. The likelihood function comes from a coalescent-based “stepping-stone” model in which discrete populations (demes) arranged on a spatial grid exchange migrants with their neighbors (Figure 1b). The parameters of this model are the migration rates between neighboring demes (*M_α,β_*) and the population sizes within each deme (*N_α_*). The prior distribution is similar to that from Petkova et al. (2016), and is based on partitioning the habitat into cells using Voronoi tesselations (one for migration and one for population size), and assuming that migration rates (or population sizes) are constant in each cell. We use an MCMC scheme to sample from the posterior distribution on the model parameters (migration rates, population sizes, and Voronoi cell configurations). We can summarize these results by surfaces showing the posterior means of demographic parameters across the habitat.

The inferred migration rates and population sizes will depend on the density of the grid used. However, using ideas from Barton et al. (2002) and Baharian et al. (2016) we convert them to corresponding parameters in continuous space, whose interpretation is independent of the grid for suitably dense grids. Specifically, we convert the migration rates to a spatial diffusion parameter *σ*(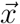), often referred to as the “root mean square dispersal distance”, which can be interpreted roughly as the expected distance an individual disperses in one generation; and we convert the population sizes (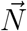) to an “effective population density” *D_e_*(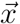) which can be interpreted as the number of individuals per square kilometer. Similar to the original grid-based demographic parameters, we can summarize MAPS results by surfaces showing the posterior means of *σ*(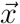) and *D_e_*(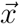) across the habitat.

### 2.2 Differences from EEMS

Our MAPS approach is closely related to the EEMS method from Petkova et al. (2016), but there are some important differences. First, the MAPS likelihood is based on lPSC sharing, rather than a simple average genetic distance across markers. This was primarily motivated by the fact that, by considering lPSC segments in different length bins, MAPS can interrogate demographic parameters across different recent time periods. However, this change also allows MAPS, in principle, to estimate absolute values for the parameters **M** and 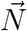, whereas EEMS can estimate only “effective” parameters which represent the combined effects of **M** and 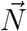. This ability of MAPS to estimate absolute values stems from its use of a known recombination map, which acts as an independent clock to calibrate the decay of PSC segments. Finally, MAPS uses a coalescent model, whereas Petkova et al. (2016) uses a resistance distance approximation (McRae, 2006).

### 2.3 Evaluation of performance under a stepping-stone coalescent model

We assess the performance of MAPS with several simulations, and compare and contrast the results with EEMS. We used the program MACS (Chen et al., 2009) to simulate data under a coalescent stepping stone model and refinedIBD (Browning and Browning, 2011, 2013) to identify lPSC segments. All simulations involved twenty demes, each containing 10,000 diploid individuals, and each exchanging migrants with their neighbors. We analyzed each simulated data set using PSC segments of length 2-6cM and *≥*6cM, which correspond to time-scales of approximately 50 generations and 12.5 generations respectively (see Lemma in the Supplementary Note). Results for other length bins are qualitatively similar (Supplementary Figure S1 & S2).

#### Migration Rate Inference

First, we simulated under a uniform (constant) migration surface with migration rate 0.01 (Figure 2a), assumed to have stayed constant over time. In this case both EEMS and MAPS correctly infer uniform migration (Figure 2a), and MAPS provides accurate estimates of the migration rate (posterior mean 0.010 when using segments 2-6cM and 0.0086 using segments *≥*6cM). As noted earlier, EEMS does not estimate the absolute migration rate; it estimates only the *relative* (effective) migration rates.

**Figure 2:**
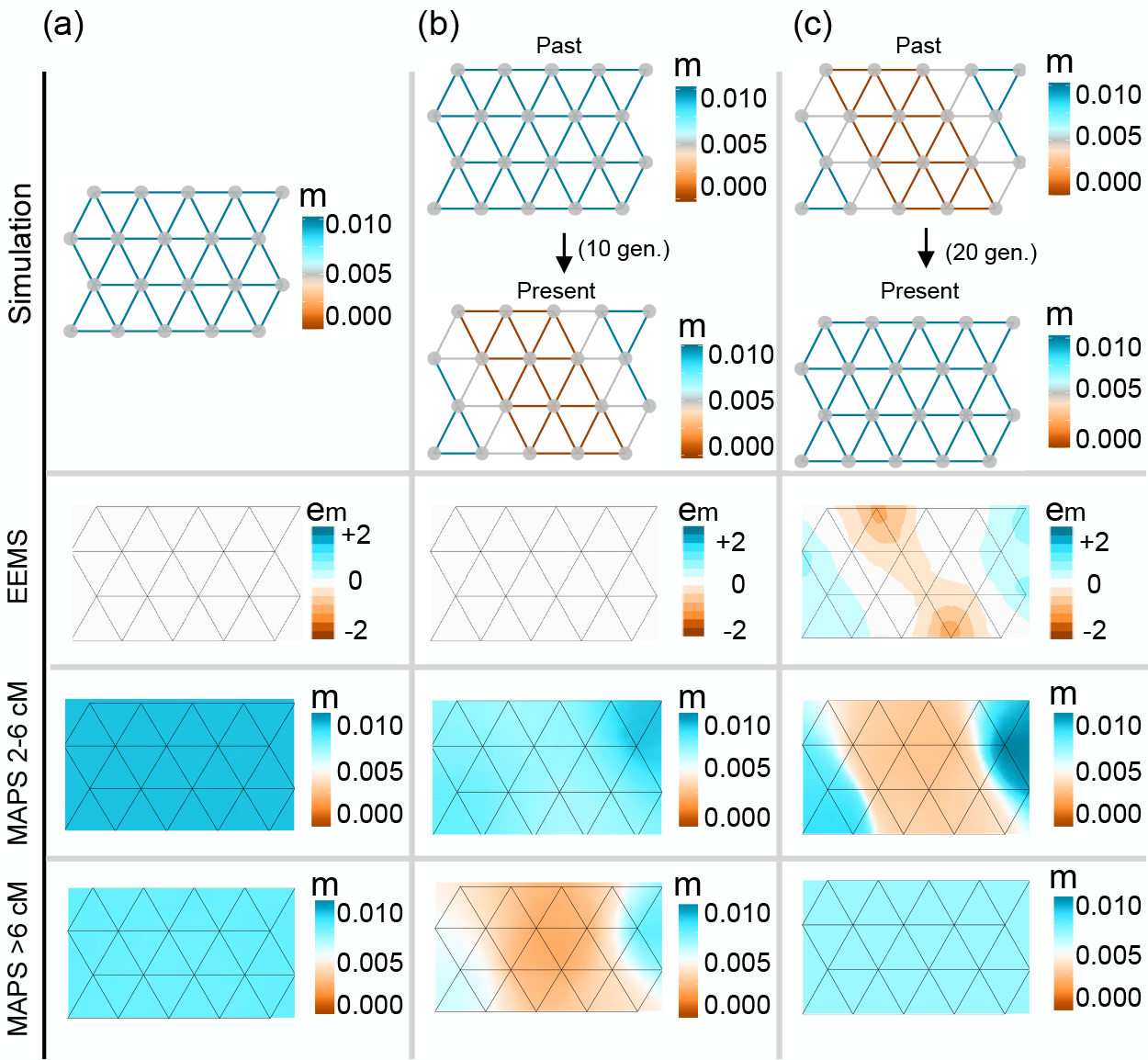
Simulations comparing migration rates inferred with MAPS against effective migration rates inferred with EEMS. (a) We simulated data under uniform migration rates equal to 0.01 and applied EEMS and MAPS using PSC segments in the range 2-6cM and *≥*6cM. Like EEMS, MAPS correctly infers a uniform migration surface. Additionally, MAPS provides accurate estimates of the migration rates for both PSC segments 2-6cM (mean 0.01) and PSC segments *≥*6cM (mean 0.0086). (b) We simulated a recent sudden migration barrier formation 10 generations ago. Here, EEMS is unable to infer a barrier, while MAPS correctly infers the historical uniform surface (2-6cM) and a barrier in the more recent time scale (*≥*6cM). (c) We simulated a long-standing migration barrier that recently dissipated 20 generations ago. EEMS infers a barrier, while MAPS correctly infers both the historical migration barrier (2-6cM) and the uniform migration surface in the more recent time scale (*≥*6cM). In all cases shown here, we simulated a 20 deme stepping stone model such that the population sizes all equal to 10,000, and 10 diploid individuals were sampled at each deme.

Next, we considered a scenario where the migration surface changed across time. Specifically the migration surface matches the constant migration scenario (above) until 10 generations ago, when a complete barrier to gene flow instantaneously arose (a “vicariance event”, Figure 2b). In this setting EEMS again infers a uniform migration surface. This is because EEMS is based on pairwise genetic distances, which are negligibly influenced by the recent barrier. In contrast, by applying MAPS with different PSC segment lengths, we can see both the historically uniform migration surface (for segments 2-6cM) and the recent barrier (segments *≥*6cM).

Next we consider a complementary time-varying scenario: an ancestral barrier disappeared 20 generations ago to allow uniform migration (Figure 2c). Here the EEMS results again reflect the longer-term processes, and a barrier is evident. And again, by applying MAPS with different PSC segment lengths, we can see different migration surfaces corresponding to different time scales, which are here reversed compared with the previous scenario: the historical barrier (for segments 2-6cM) and the recent uniform migration (segments *≥*6cM).

#### Population Size Inference

As noted above, and discussed in (Petkova et al., 2016), EEMS estimates an “effective” migration surface that reflects the combined effects of population sizes 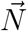 and migration rates **M**; consequently it cannot distinguish between variation in **M** and variation in 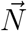. In contrast, MAPS has the potential to distinguish these two types of variation.

To illustrate this difference we simulate data with a constant migration surface, and a population size surface that has a 10-fold “dip” in the middle of the habitat (deme size 1,000 vs 10,000; Figure 3). Petkova et al. (2016) performed a similar simulation, and showed that EEMS estimated an effective migration surface with an “effective barrier” in the middle, caused by the dip in population size. As expected, we obtain a similar result for EEMS here. Further, the EEMS inferred diversity surface is also approximately constant, because the diversity surface reflects changes in within-deme heterozygosity, and these vary little in this simulation. In contrast, MAPS is able to separate the influence of migration and population sizes: the estimated migration surface is approximately constant (with mean migration rate equal to the true value 0.01) and the estimated population size surface shows a dip in the middle, with accurate estimates of deme sizes (mean 985 at the center and 9100 at the edges). Additional simulations with non-uniform migration rates reinforce these results; see Supplementary Figure S3.

**Figure 3:**
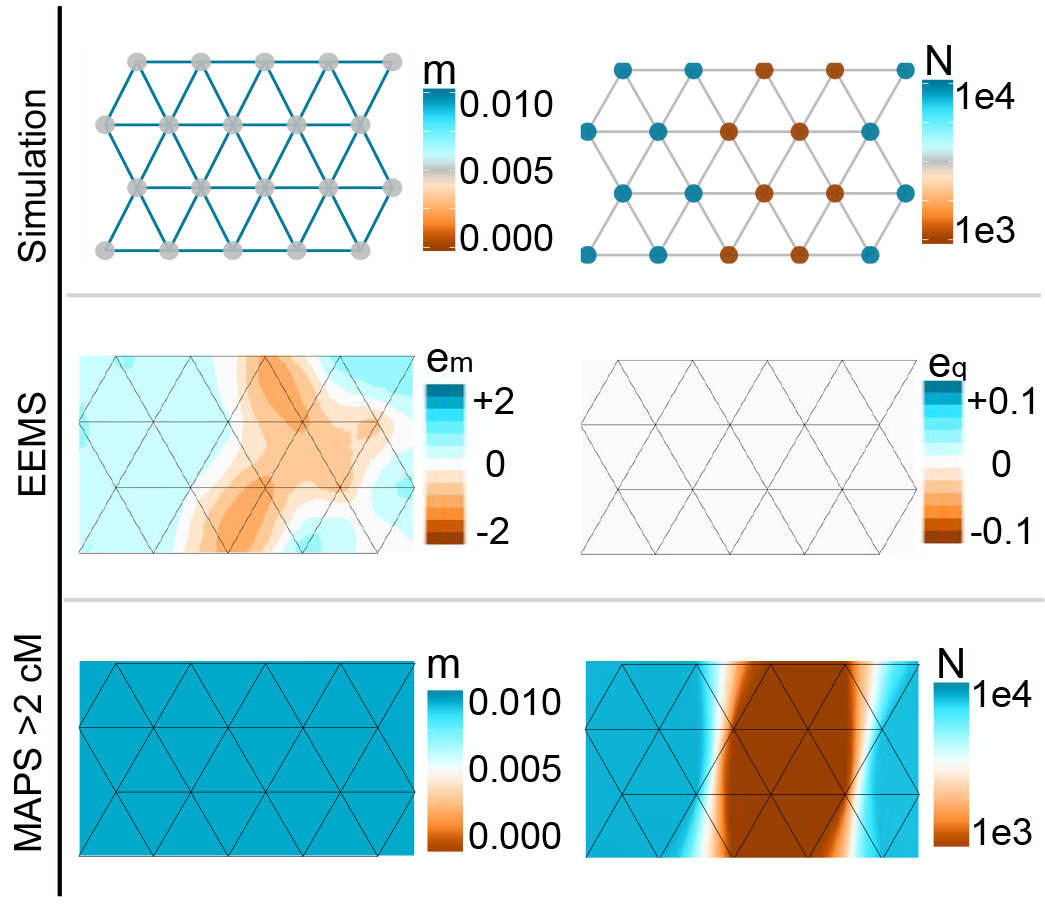
Simulations comparing population sizes inferred with MAPS and “diversity-rates” inferred with EEMS. We simulated uniform migration rates of 0.01 and a trough of low population sizes in the center of the habitat such that population sizes equal to 1,000 at the center and 10,000 otherwise. Under these simulations, EEMS infers a barrier in effective migration and infers uniform diversity rates. However, MAPS correctly infers a uniform migration surface (mean 0.01) and provides accurate estimates of deme sizes (mean 985 at the center and 9100 at the edges)

### 2.4 Applying MAPS to the POPRES data

To illustrate MAPS on real data, we analyze a genome-wide SNP dataset on individuals of European ancestry (the “POPRES” study Nelson et al., 2008). Previous analyses of these data have shown the strong influence of geography on patterns of genetic similarity (Novembre et al., 2008; Lao et al., 2008; Ralph and Coop, 2013). In particular Ralph and Coop (2013) analyzed spatial patterns in the sharing of PSC segments across Europe. To facilitate comparison with their results, we use their PSC segment calls, focusing on a subset of 2224 individuals after filtering (see Methods).

We applied MAPS to these data using three different PSC segment length bins: 1 *−* 5cM, 5 *−* 10cM, and > 10cM. The longer bins correspond to more recent demography because as PSC lengths increase, the average coalescent times decrease. Indeed, the average coalescent times for each of these three length bins is inferred to be 90, 23 and 7.5 generations respectively (Supplementary Note), which correspond to 2700 years, 675 years and 225 years if we assume 30 years per generation.

We note that the accuracy of called PSC segments will vary across these bins: based on simulations in Ralph and Coop (2013) PSC segment calls in the smallest bin (1-5cM) will likely suffer from both false positives and false negatives, whereas for the longer bins PSC calls should be very reliable. Nonetheless, even in the smallest bin, closely-related individuals will still tend to show higher PSC sharing, and so the estimated MAPS surfaces should provide a useful qualitative summary of spatial patterns of variation even if quantitative estimates may be less reliable.

#### Inferring dispersal and population density surfaces

The inferred MAPS dispersal rates (migration rates scaled by grid step size) and population densities (population sizes scaled by grid area size) for each PSC length bin are shown in Figure 4.

**Figure 4:**
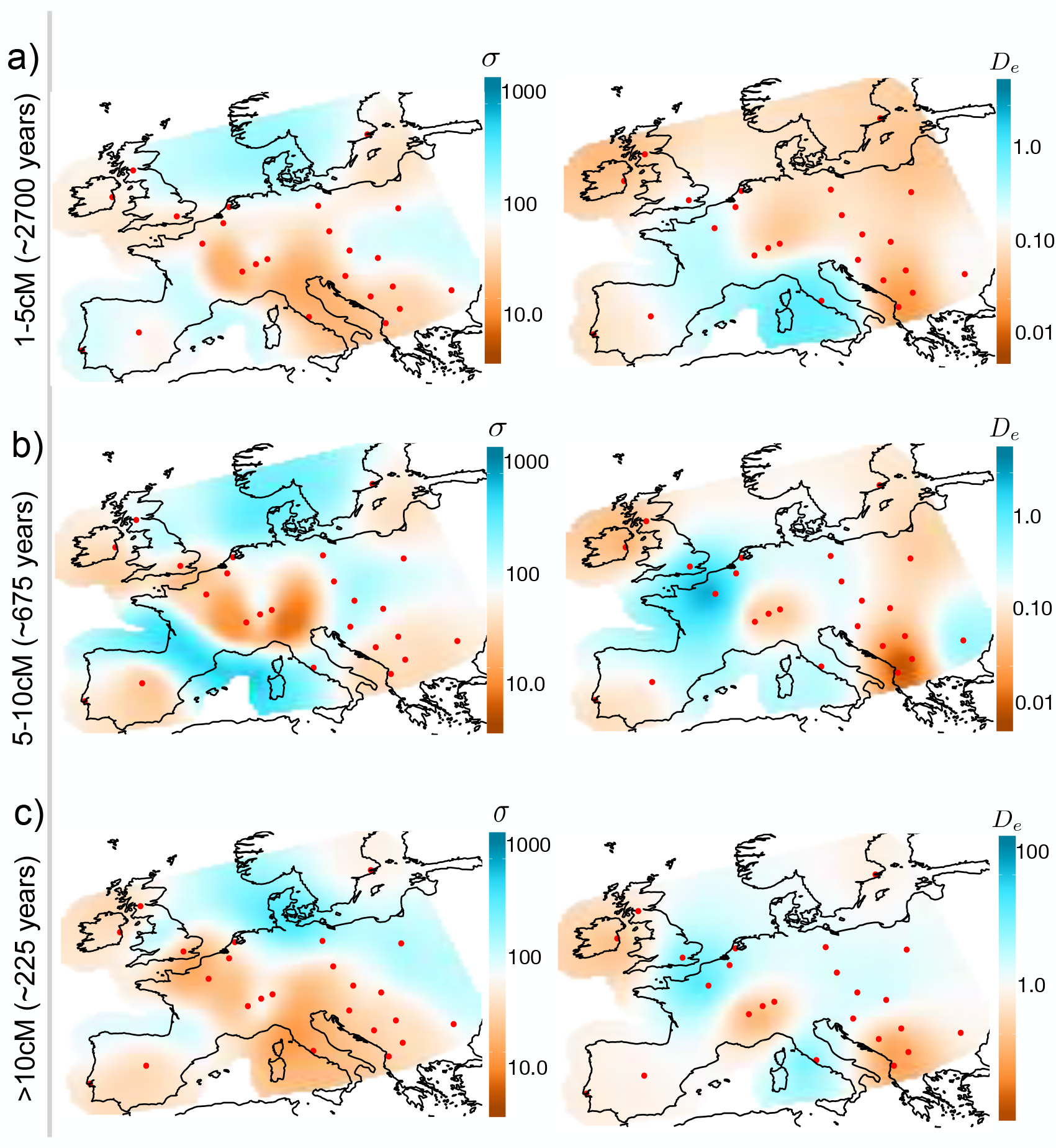
Inferred Dispersal Surfaces and Population Density Surfaces over time for Europe. We apply MAPS to a European subset of POPRES Nelson et al. (2008) with 2,234 individuals and plot the inferred dispersal *σ*(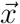) and population density *D_e_*(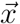) surfaces for PSC length bins (a) > 1cM (b) 5-10cM and (c) >10cM. We transform estimates of 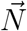 and **M** to estimates of *σ*(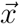) and *D_e_*(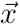) by scaling the migration rates and population sizes by the grid step-size and area (see Equations (17) and (18)). Generally, we observe the patterns of dispersal to be relatively constant over time periods, however, we see a sharp increase in population density in the most recent time scale (>10cM). Note the wider plotting limits in inferred densities in the most recent time scale.

Largely speaking, the spatial variation in inferred dispersal rates and population densities is remarkably consistent across the different time scales (Figure 4). In the MAPS dispersal surfaces, several regions with consistently low estimated dispersal rates coincide with geographic features that would be expected to reduce gene flow, including the English Channel, Adriatic Sea and the Alps. In addition we see consistently high dispersal across the region between the UK and Norway, which may reflect the known genetic effects of the Viking expansion (e.g Leslie et al., 2015). The MAPS population density surfaces consistently show lowest density in Ireland, Switzerland, Iberia, and the southwest region of the Balkans. This is consistent with samples within each of these areas having among the highest PSC segment sharing (Supplementary Figure S4a). The MAPS inferred country population sizes are also highly correlated with estimated current census population sizes from The World Bank (2016) and National Records of Scotland (2011) (Supplementary Figure S6).

The most notable variation among the estimated surfaces from different time scales is a dramatic increase in the mean estimated population density in the most recent time scale (Figure 4 and Supplementary Figure S7). Indeed, the estimated mean for the last time scale – 1.4 individuals per square km – is 6-9 fold higher than those for the earlier time scales (0.16 and 0.22 respectively). This increase is consistent with the recent exponential growth of human population sizes (Cohen, 1995). The estimates themselves are lower than historical estimates of *≈*1-30 individuals per square km based on archaeological data (e.g. Zimmermann et al., 2009).

The dispersal surfaces show more minor changes between time periods (Figure 4 and Supplementary Figure S7). In particular, the estimated mean dispersal rates are relatively constant across time, being 73, 103 and 72 respectively (in units of km in a single generation). These mean estimates are consistent with empirical estimates of 10-100 km in a single generation compiled by Kaplanis et al. (2018) using pedigrees of individuals living between 1650 and 1950 AD. We do note the lower estimated dispersal rates between Portugal and Spain in the analyses of longer PSC segments (5-10 and > 10cM), and the higher estimated dispersal rates through the Baltic Sea (> 10cM segments), possibly reflecting changing gene flow in these regions in recent history.

#### Comparison to Ringbauer et al. (2017)

Ringbauer et al. (2017) also estimate a mean dispersal rate and population density from the Eastern European subset of the data analyzed here. Their estimates are based on PSC segments > 4cM, which is most comparable with our analysis of 5-10cM. Unlike our analysis, their estimates are based on a spatially homogeneous model. To compare with their estimates we computed the mean of the estimated dispersal rate and population densities in Eastern Europe (but based on an analysis of the full data). For the dispersal rate this yields an estimate of 88 km in a single generation, which is consistent with the range of 50-100 given by (Ringbauer et al., 2017). For the population density, it yields an estimate of 0.10 individuals per square km, which is somewhat higher than the estimate of 0.05 obtained under a comparable (time-homogeneous) population model in (Ringbauer et al., 2017). Possibly our higher estimate partly reflects the influence of our spatial modeling approach, which will tend to shift the estimate for Eastern Europe toward the estimated mean across all of Europe (which is 0.22). In addition, the difference in length thresholds (> 4cM versus 5-10cM) may also be contributing; if segments in the Ringbauer et al. (2017) analysis are on average shorter and hence older, one would expect lower density estimates, based on our results that suggest lower densities in the past (Figure 4).

#### Comparison with EEMS

The EEMS results for these data (Figure S8) show non-trivial differences with the MAPS results (Figure 4a). Two potential causes are: i) differences in the summary data used (PSC segment sharing vs genetic distances) and hence sensitivity to different timescales; and ii) differences in the underlying models (e.g. composite Poisson likelihood vs Wishart likelihood, and different parameterizations/approximations to the coalescent model; see Discussion). To evaluate the impact of i) we compared the PSC segment sharing and genetic distances, and found their correlation to be only modest (Pearson’s *ρ* = −0.38), with the most notable deviation for comparisons between countries in Eastern Europe (Figure S9a). Furthermore, most of this correlation is due to geographic distance: after controlling for geographic distance the correlation is only −0.18, which may be a more relevant metric because inferred spatial heterogeneity in gene flow (barriers and corridors) is driven by departures from simple isolation by distance.

To better assess the impact of ii) we applied EEMS on a distance matrix constructed to have the same similarity patterns as the PSC segment sharing matrix input to MAPS (1*−*5cM length bin). The resulting EEMS surface is more similar to the corresponding MAPS dispersal surface (Supplementary Figure S9b vs Figure 4a), but there remain substantial differences. This investigation confirms what we expected *a priori* — the two surfaces should be different because the underlying models and inferred parameters of MAPS and EEMS are different. As noted before, EEMS infers the “effective migration rate” which reflects the effects of both the migration rates and population sizes, while MAPS infers them separately.

## 3 Discussion

We developed a method (MAPS) for inferring migration rates and population sizes across space and time periods from geo-referenced samples. Our method builds upon a previous method developed for estimating effective migration surfaces (EEMS) (Petkova et al., 2016). However there are several differences between MAPS and EEMS. Most fundamentally, MAPS draws inferences from observed levels of PSC sharing between samples, whereas EEMS draws inferences from the genetic distance. These two data summaries capture different information about the coalescent distributions: in essence, PSC sharing captures the frequency of recent coalescent events, whereas genetic distance captures the mean coalescent time. Consequently MAPS inferences largely reflect the recent past (⪅ 1000 years for human recombination rates and generation times with PSC segments > 2cM), whereas EEMS inferences reflect demographic history on a longer timescale across which pairwise coalescence occurs (99% of events > 6000 years old, assuming diploid *N_e_* of 10,000 for humans, exponential coalescent time distribution).

Another consequence of modelling PSC sharing, rather than genetic distance, is that MAPS can separately estimate demographic parameters related to migration rates (**M**) and population sizes (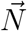), as in Figure 3 for example. In essence MAPS does this by using the known recombination map as an additional piece of information to help calibrate inferences. In contrast EEMS, which makes no use of recombination maps, cannot separate **M** and 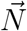. Instead EEMS infers a compound parameter referred to as the “effective migration rate”, which is influenced by changes in both **M** and 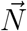; see Figure 3. In principle, if applied to sequence data instead of genotype data at ascertained SNPs, the genetic distances used by EEMS could perhaps also separately estimate **M** and 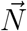 by exploiting known mutation rates to calibrate inferences. However, this would require non-trivial additional changes to the current EEMS likelihood, which was designed to be applicable to ascertained SNPs and does not explicitly model variation in population sizes. (The EEMS likelihood instead uses a “diversity rate” *e_q_*, which reflects within-deme heterozygosity but is not explicitly a population size parameter.)

An additional useful feature of PSC segments is that, by varying the lengths analyzed, one can infer parameter values across different time scales. For example, our simulations show how by contrasting shorter and longer PSC segments, the method can reveal different gene flow patterns in scenarios with recent changes (see Figures 2 and 3). Further support comes from our empirical analysis of the POPRES data-set, where we found population sizes inferred from longer PSC segments to be more correlated with census sizes The World Bank (2015 census 2016) and National Records of Scotland (2011, 2011 census) than sizes inferred from shorter segments (e.g. Spearman’s *ρ* = 0.71 for 1 *−* 5cM and *ρ* = 0.84 for > 10cM; see Supplementary Figures S5 and S6). Also, not surprisingly, PSC segments greatly outperform using heterozygosity as an indicator of census population size (the Spearman’s correlation coefficient between heterozygosity and census size was insignificant, p-value = 0.25).

Our estimates of dispersal distances and population density from the POPRES data are among the first such estimates using a spatial model for Europe (though see (Ringbauer et al., 2017)). The features observed in the dispersal and population density surfaces are in principle discernible by careful inspection of the numbers of shared PSC segments between pairs of countries (e.g. using average pairwise numbers of shared segments, Supplementary Figure S4b, as in Ralph and Coop (2013)). For example, high connectivity across the North Sea is reflected in the raw PSC calls: samples from the British Isles share a relatively high number of PSC segments with those from Sweden (Supplementary Figure S4b). Also the low estimated dispersal between Switzerland and Italy is consistent with Swiss samples sharing relatively few PSC segments with Italians given their close proximity (Supplementary Figure S4b). However, identifying interesting patterns directly from the PSC segment sharing data is not straightforward, and one goal of MAPS (and EEMS) is to produce visualizations that point to patterns in the data that suggest deviations from simple isolation by distance.

Our results suggest that several features of dispersal in Europe have been relatively stable over the last *~*3000 years, whereas the population sizes have been increasing. The relative stability of the gene flow patterns is perhaps surprising given ancient DNA results suggest a continually dynamic history of population movements. One possibility is that much of European population structure may have been established by the end of the Bronze Age (4,000 years ago), with relatively more stable patterns in the intervening period that is reflected in lPSC segments. Nonetheless, the dispersal is not completely stable– our results suggest changes in Iberia, the Baltic, and to minor degrees in other areas.

The inferred population size surfaces for the POPRES data show a general increase in sizes through time, with small fluctuations across geography; for instance, Polish samples have a relatively larger population size in inferred values from the largest length scale (> 10cM). In our results, the smallest inferred population sizes are in the Balkans and Eastern Europe more generally. This is in agreement with the signal seen by Ralph and Coop (2013); however, taken at face value, our results suggest that high PSC sharing in these regions may be due more to consistently low population densities than to historical expansions (such as the Slavic or Hunnic expansions).

Although consistent with previous results, our estimates of dispersal and population sizes do not exactly agree with empirical estimates. For example, our estimates of population sizes are consistently lower than the census sizes (Supplementary Figure S6). This is to be expected for several reasons. First, census sizes include non-breeding individuals (juvenile and post-reproductive age) that do not impact the formation of PSC segments. Second, MAPS is fitting a single population size per location, and in a growing population the best fit population size will be an under-estimate of contemporary size. Third, in a wide class of population genetic models, the effective size, even among reproductive age individuals, is lower than the census size because of factors that inflate the variance in offspring number. Fourth, some discrepancy is expected simply because the stepping-stone population genetic model used here is only a coarse approximation to the complex spatial dynamics of human populations. Finally, recombination rate mis-specification can bias the inferred parameters. Furthermore, we caution that our results must be interpreted in the light of the fact that we have limited spatial sampling across Europe, and only very coarse geographical origin data (country of origin).

Here, as in Petkova et al. (2016) we use a discrete stepping-stone model to approximate a process that might be more naturally modelled as continuously varying in space. Recent work (Ringbauer et al., 2017; Baharian et al., 2016) exploits continuous models to estimate dispersal and population density parameters from sharing of lPSC segments. However, these methods assume that dispersal and population density are constant across space: extending them to allow these parameters to vary across space could be an interesting avenue for future work.

A major achievement in method development in population genetics would be to jointly infer migration rates and population sizes across both space and time. MAPS is a step towards this goal. However, we do not infer demography explicitly as a function of time and instead infer surfaces in time blocks defined by PSC length bins. In principle, our method allows for inference of demography across time by treating PSC segments as independent across length bins, see Equation (S27) in Supplementary Methods. However, this requires fitting multiple migration/population surfaces and is computationally unfeasible with our current MCMC routine. Other computational techniques (e.g. Variational Bayes or fast optimization of the likelihood) might make this goal possible.

## 4 Methods

### 4.1 MAPS configuration

For the empirical data analysis, we ran MAPS with 200 demes. The MAPS output was obtained by averaging over 20 independent replicates (the number of MCMC iterations in each replicate was to set 5e6, number of burn-in iterations set to 2e6, and we thinned every 2000 iterations). We provide the the MAPS here: https://github.com/halasadi/MAPS, and the plotting scripts here: https://github.com/halasadi/plotmaps.

### 4.2 Inferring PSC segments from the data

Our pipeline to call PSC segments for simulations can be found here: https://github.com/halasadi/ibd_data_pipeline. We follow the recommendations of Browning and Browning (2011, 2013) and Ralph and Coop (2013) by running BEAGLE multiple times and merging shorter segments.

For the empirical data analysis, we use the PSC segments (“IBD”) calls from Ralph and Coop (2013), which can be found here: https://github.com/petrelharp/euroibd. We further applied a filter to retain countries with at least 5 sampled individuals, and removed Russian and Greek individuals to restrict the habitat to a smaller spatial scale

### 4.3 Model

MAPS assumes a population genetic model consisting of triangular grid of *d* demes (or populations) with symmetric migration. The density of the grid is pre-specified by the user with the consideration that the computational complexity is *O*(*d*^3^). We use Bayesian inference to estimate the MAPS parameters: the migration rates and coalescent rates *M* and *<u>q</u>* respectively. Its key components are the likelihood, which measures how well the parameters explain the observed data, and the prior, which captures the expectation that *M* and *<u>q</u>* have some spatial structure (in particular, the idea that nearby edges will tend to have similar migration rates and nearby demes have similar coalescent rates).

MAPS estimate the posterior distribution of Θ = *M, <u>q</u>* given the data. The data used for MAPS consists of a similarity matrix 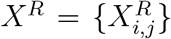 which denotes the number of PSC segments in a range *R* = [*u, v*] base-pairs shared between pairs of haploid individuals (*i, j*) ∈ {1, ⋯, *n*} × {1, ⋯, *n*} where *n* is the number of (haploid) individuals. Furthermore, a recombination rate map is required as input for MAPS. The likelihood is a function of the expected value of 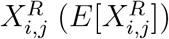. Below we describe the computation of 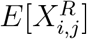 and the other key components of the likelihood. Finally, we briefly describe the prior used and an MCMC scheme to sample from the posterior distribution of Θ.

#### The likelihood function

Let *α, β* denote the demes that (haploid) individuals *i* and *j* are sampled in, we define,

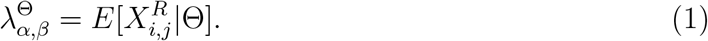

For the marginal distribution, we assume

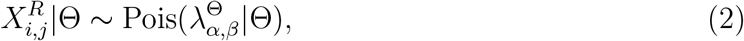

and one option for computing the joint distribution of the data is to assume independence between pairs of individuals (*i, j*) as done previously (Palamara et al., 2012; Palamara and Peer, 2013; Ralph and Coop, 2013; Ringbauer et al., 2017). This assumption leads to the log-likelihood,

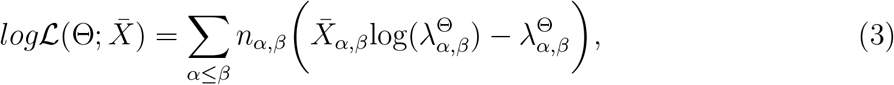

where *X* = {*X*_*α,β*_} such that (*α, β*) ∈ {1, ⋯, *d*} × {1, ⋯, *d*} and *d* is the number of demes. Furthermore

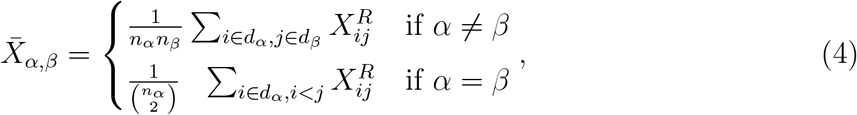

where *n_α_* is the number of sampled individuals in deme *α*, *d_α_* is the set of all individuals in deme *α*, and

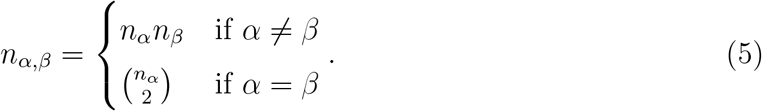

However, we found that there were significant correlations in lPSC segments between individuals. To deal with this, we down-weighted the likelihood function to reflect the “effective” number of samples (*e_α,β_*) instead of the number of pairs (*n_α,β_*). The effective number of samples between demes *α,β* is given by,

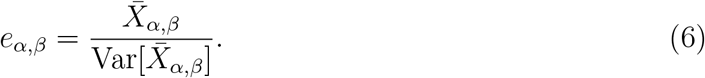

In the case of independence, 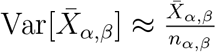. However, because of correlations in the data, the actual variance is significantly larger than the variance computed under an independence model. Here, we estimate Var[*X_α,β_*] by bootstrapping individuals with replacement. This way, we model the correlations between pairs of individuals for within and between-deme comparisons. The loglikelihood adjusted for correlations is given by,

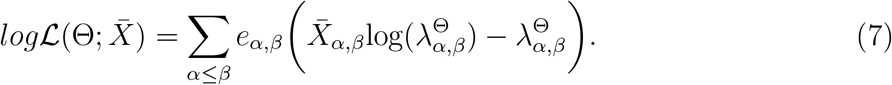

#### Computing the expectation of 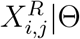

Next, we derive expressions to compute the expectation of the number of PSC segments of length greater than 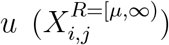 conditional on the demography Θ. From results in Palamara et al. (2012) it is easy to show that

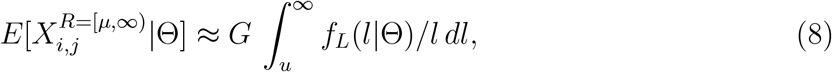

where *G* denotes the length of the genome (in base-pairs), *L* denotes the random length (in base-pairs) of the PSC segment between *i* and *j* containing a pre-specified position in the genome (base *b* say), and *f_L_* is its probability density. Intuitively, *Gf_L_*(*l|*Θ) is the expected number of base-pairs that lie in PSC segments of length *l*, making 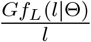 the expected number of PSC segments of length *l*. Integrating the latter quantity from *µ* to *∞* gives the desired result.

To help compute (8) we introduce *T_ij_* to denote the (random) coalescent time in generations between *i* and *j* at base *b*, with density *f_T_ij__* (*t|*Θ). Then (8) can be written as an integral over *T_ij_*:

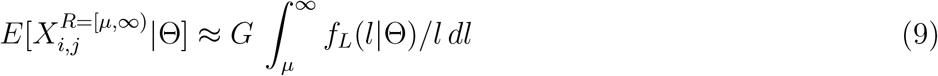

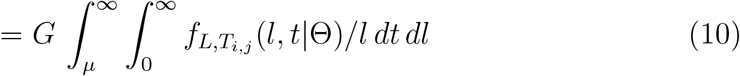

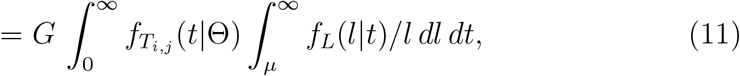

using the relation that *f_L,T_i,j__* (*l, t|*Θ) = *f_L_*(*l|t*, Θ)*f_T_i,j__* (*t|*Θ) = *f_L_*(*l|t*)*f_T_i,j__* (*t|*Θ). A key simplification here comes from the fact that, given *T_ij_*, *L* is conditionally independent of Θ.

It can be shown that the conditional distribution of *L* given *T_ij_* is an erlang-2 distribution (Palamara et al., 2012; Palamara and Peer, 2013; Hein et al., 2004) with density

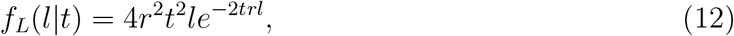

where *r* is the recombination rate per base-pair. Substituting this into the inner integral of (11) and integrating analytically yields

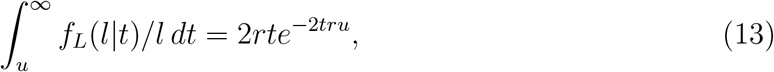

leading to

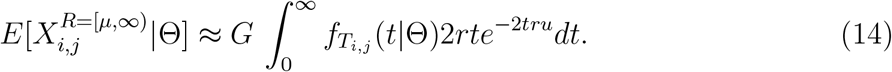

Here, we assume the probability density of *T_i,j_* is given by,

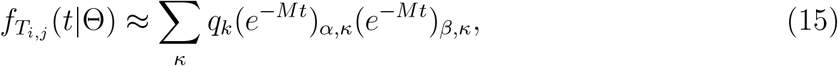

where demes *α, β* denote the deme where lineages *i* and *j* are sampled from, 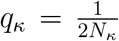 is the coalescent rate in deme *κ*, and *M* = *〈m_α,β_〉* is the migration rate matrix between all *d* demes such that (*α, β*) ∈ {1, ⋯, *d*} × {1, …, *d*}. We compute the matrix exponential by first diagonalizing the matrix *M* = *P DP^T^* and compute *e^−M t^* = *P e^−Dt^P^T^*.

Having computed all individual components of 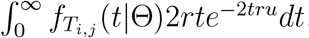, we are left to evaluate a one-dimensional integral which we do by Gaussian quadrature (with 50 weights).

To compute the expected number of PSC segments in a range *R* = (*µ, ν*)

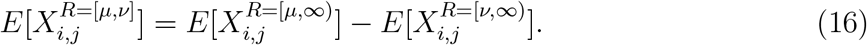

As mentioned previously, the units of *µ, ν* are in base-pairs. However, we can transform to units of centiMorgans (cM) by: *µ_cM_* = 100*µr*.

#### The Prior

MAPS uses a hierarchical prior parameterized by Voronoi tessellation (similar to EEMS). The Voronoi tessellation partitions the habitat into *C* cells. Given a Voronoi tessellation of the habitat, each cell *c* ∈ {1, ⋯, *C*} is associated with a migration rate (*ℳ_c_*) and population size (*𝒩_c_*). Demes (*α*) that fall into cell *c* will have population size *N_α_* = *𝒩_c_*, and similarly, migration rates between demes *α, β* equal 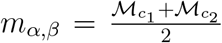 if demes *α, β* fall into cells *c*_1_ and *c*_2_. We use an MCMC to integrate over the distribution on partitions of Voronoi cells. See Supplementary Notes section 5.4 for more information.

#### The MCMC

We break up the MCMC updates into updating a series of conditionally independent distributions. Provided the conditional posterior distributions for each part give support to all the parameter space, this will define an irreducible Markov chain with the correct joint posterior distribution Stephens (2000). We use Metropolis-Hastings to update all parameters, and random-walk proposals for most updates, with exception of birth-death updates for updating the number of Voronoi cells. See Supplementary Notes section 5.5 for more information.

#### Transformation of parameters to continuous space

Given an inferred population size at a particular deme *α* and a grid with uniform spacing, the transformation from population size to population density is given by

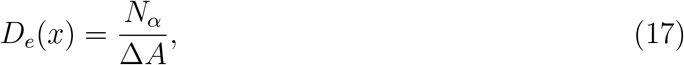

where 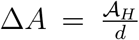 is the area covered per deme such that 풜_*H*_ is the area of the habitat (in km^2^), *d* is the number of demes, and *x* corresponds to the spatial position of deme *α*. Intuitively, (17) implies that the density multiplied by the area equals population size, i.e. *D_e_*(*x*)∆*A ≈ N_α_*. Equation (17) can is analogous to equation 7 in (Baharian et al., 2016).

Given a migration rate (*m*), the transformation to dispersal distances is given by,

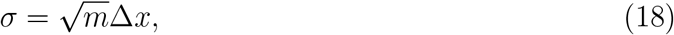

where ∆*x* is the step size of the grid (km). The dispersal distance represents the distance traveled by an individual after one generation, and sometimes is called the “root mean square distance” or “dispersal rate” (Barton et al., 2002). Please see Supplementary Note section 5.2 for the derivation of (18).

## Acknowledgements

We thank Dick Hudson for helpful discussion on the identifiability of demographic parameters in coalescent models; Evan Koch, Ben Peter and the Novembre & Stephens Lab for comments on the paper and helpful discussion. We further thank Peter Carbonetto for computational support and helpful discussion.

## 5 Supplementary Note

### 5.1 The model

The coalescent process for two samples under a multi-deme model can be described by a continuous time Markov chain (CTMC) (Bahlo and Griffiths, 2001). Let *i, j* represent sampled lineages and *α, β* their locations, respectively, *d* is the number of demes (or populations) and (*α, β*) ∈ {1, ⋯, *d*} × {1, …, *d*}. Let *c* denote the coalescent state. The infinitesimal rate matrix *R* of this CTMC is

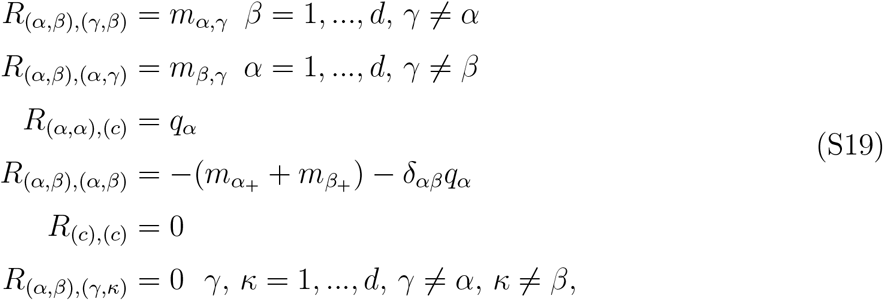

where *M* = *〈m_α,β_〉* denotes the migration rate matrix, and *m_α,β_* is the migration rate between demes *α, β* and 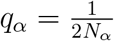 is the coalescent rate of deme *α* which is proportional to the inverse of the population size at deme *α* (*N_k_*). Let *T_i,j_* denote the (random) coalescent time between the pair of sampled lineages, and *f_T_i,j__* (*t*) denote the probability density of a coalescent event at time *t*. Here, we derive *f_T_i,j__* (*t*) by conditioning on the position of the two lineages.

#### Lemma 5.1

*Let X_i_*(*t*), *X_j_* (*t*) ∈ {1, ⋯, *d*} × {1, …, *d*} *denote the position of lineage i and lineage j at time t respectively. The probability density f_T_i,j__ (t) that lineage i and j coalesce at time t is given by 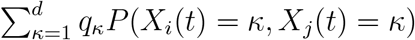*

For ∆*t ≈* 0,

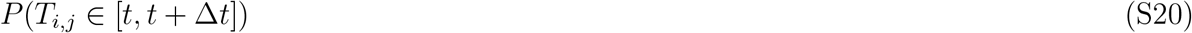

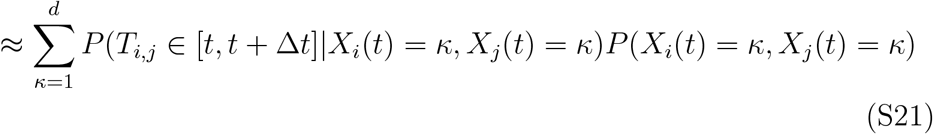

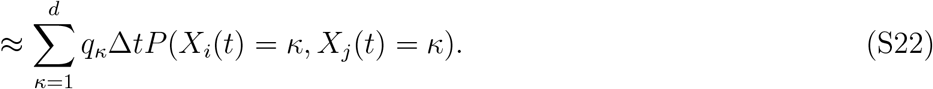

Taking the limit ∆*t →* 0, we arrive at the density

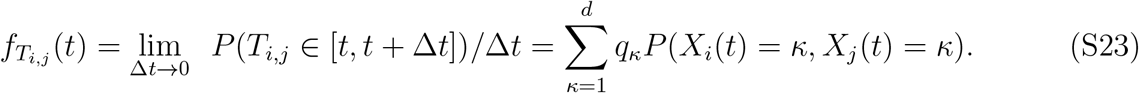

#### The random walk approximation to the coalescent

Here, we introduce an approximation,

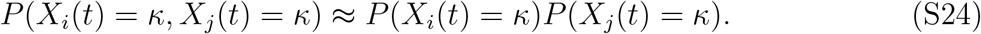

The intuition is that probability that lineage *i* and *j* coalesce before time *t* is extremely small such that the two lineages approximately behave like two independently moving particles. Each lineage can be modeled by a random walk with transition matrix *M*. These assumptions were also made in the context of continuous spatial diffusion models for haplotype sharing Baharian et al. (2016); Ringbauer et al. (2017), and even further back, as a general approximation to the two-dimensional continuous-space coalescent process (Barton et al., 2002; Wilkins, 2004; Blum et al., 2004; Novembre and Slatkin, 2009; Robledo-Arnuncio and Rousset, 2010).

This approximation implies that

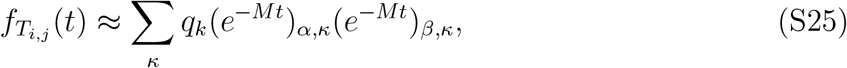

where lineages *i, j* are initially sampled in deme *α, β*. Or equivalently in matrix form,

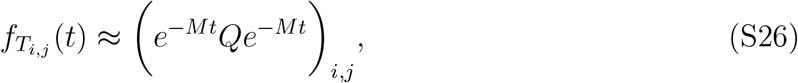

where *Q* = *diag*(*q*_1_, …, *q_d_*).

#### Varying migration rates and population sizes across time

##### Corollary 5.1.1

*Let time slice k be defined by the interval t_k−_*_1_ *< t < t_k_, M_k_ denote the migration rate matrix in time slice k, and 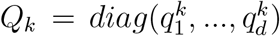 where 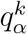 denotes the coalescent rate in deme α at time slice k. Let T_i,j_ denote the coalescent time between lineage i, j sampled in demes α, β, then under the independence assumption, for t ∈* (*t_K−_*_1_, *t_K_*),

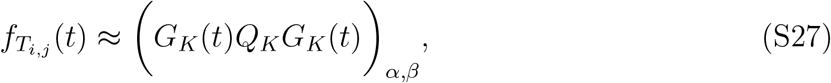

*where 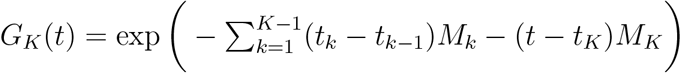*.

#### Expected number of lPSC segments given the demography Θ

##### Lemma 5.2

*Let 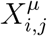 denote the number of PSC segment greater than µ basepairs shared between haploid individuals i, j*, Θ *denote the demographic model, G the size of the genome, L denotes the random length (in base-pairs) of the PSC segment between i and j containing a pre-specified position in the genome, then 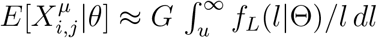*.

Let *E*[*ℱ^μ^|*Θ] denote the expected fraction of the genome between *i, j* that lies in PSC segments greater than *μ*, and *E*[*s^μ^|*Θ] the expected size of a PSC segment conditional on it being at least length *μ*. According to equations 9-14 from (Palamara et al., 2012),

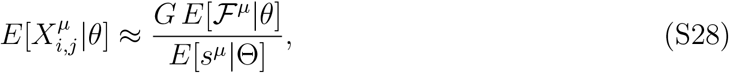

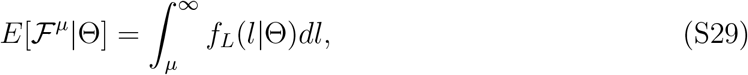

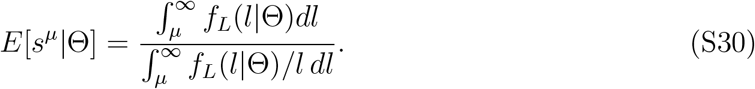

We obtain the desired result by substituting (S29) and (S30) into (S28) and canceling liketerms.

#### Expected age of a segment

We choose PSC segment lengths based on their expected age which is derived below.

##### Lemma 5.3

*The expected coalescent time (t, in generations) of an PSC segment between between length L*_1_ *centiMorgans and L*_2_ *centiMorgans is approximately 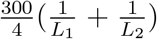 if the effective population size (N) is sufficiently large*.

We choose to work in units of basepairs, and will convert back to units of morgans at the end. We convert *L*_1_ into units of base-pairs with the transformation: 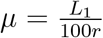 and similarly 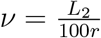

Let us denote *T |l, N* as the random coalescent time of a PSC segment that is at least length *l* under a single-deme demography model with population size *N*. The expected coalescent time of an PSC segment longer than *μ* base-pairs can be expressed as

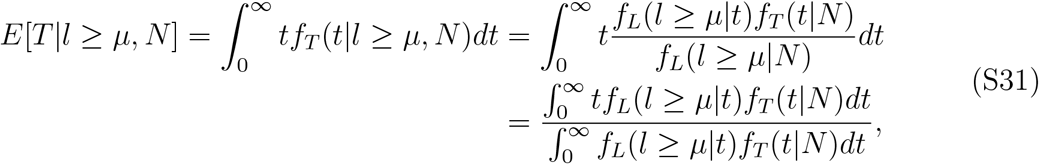

where *f_L_*(*l|t*) = 4*r*^2^*t*^2^*le*^−2*trl*^ denotes the probability density that a PSC segment is of length *l* given it has a common ancestor event at time *t*, *f_T_* (*t|N*) denotes the probability density that a coalescent event occurs at time *t* under the demography model with population size *N*.

Next, we expand a key term in equation (S31)

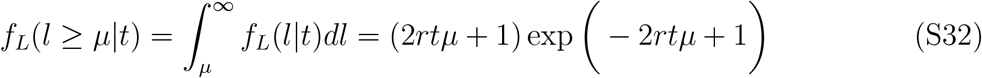

and assume,

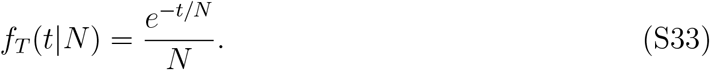

Putting everything together,

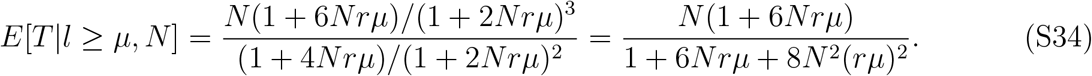

We can remove the dependence of *N* by taking lim_*N →∞*_ as done similarly in Baharian et al. (2016),

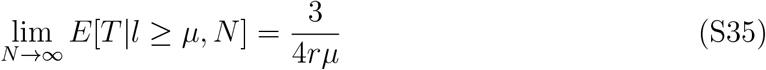

Now that we have derived the expected age of PSC segment longer than *μ*, it is quite simple to expand the equation for PSC segments between *μ* and *ν* base-pairs,

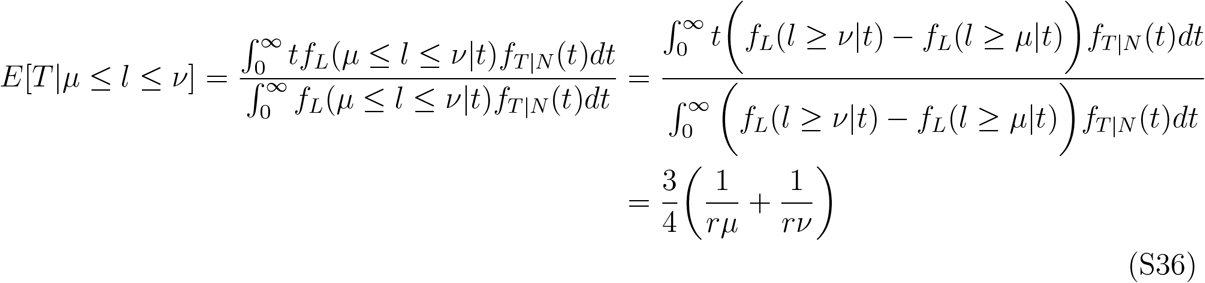

We transform back to units of centimorgans: let *L*_1_ = 100*rμ* and *L*_2_ = 100*rν* be in units of centiMograns, and taking the limit, we get the desired result

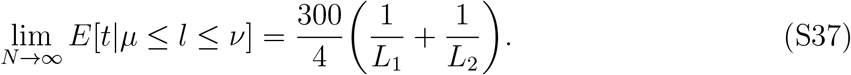

### 5.2 Transformation of migration rates to dispersal rates

Migration rates inferred under a discrete model can be transformed to dispersal distances representing parameters in continuous space. Here, we derive the transformation.

#### Lemma 5.4

*Consider a random walk on a 2D grid, where steps are taken according to a Poisson process with rate m, and let <u>X</u>*(*t*) *be a vector denoting the coordinates of the particle at time t. The distribution of <u>X</u>*(*t*) *approximately only depends on the compound parameter m*(∆*x*)^2^ *(or equivalently* 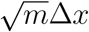).

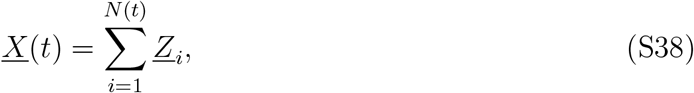

where *N* (*t*) is the number of steps taken by time *t*, and *<u>Z</u>_i_* is a random variable representing the direction and magnitude taken at step *i*. Since *X*(*t*) is a sum of iid variables, a form of the central limit theorem applies here and *X*(*t*) converges to the normal distribution (Rényi, 1960).

In a random walk on a triangular grid, a particle can move in one of the 6 directions (upper-right, right, lower-right, left, upper-left, and lower-left):

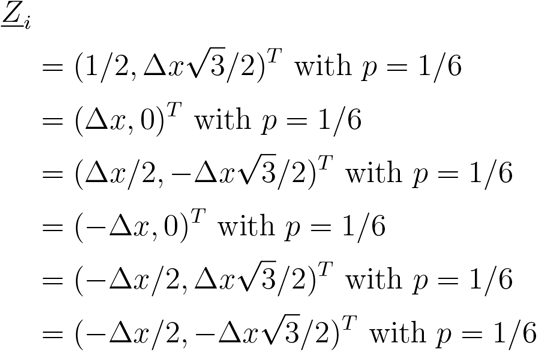

where ∆*x* represents the step size in the grid (i.e. edge length). The mean and variance are given by,

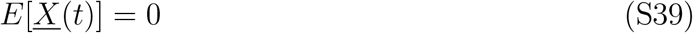

and,

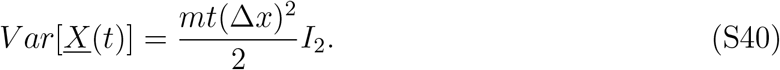

where *I*_2_ is the identity matrix. Under normality, the mean and variance are sufficient statistics. Note that (S39) and (S40) also hold for square grids.

#### Interpretation of the migration diffusion parameter *m*(∆*x*)^2^

In addition, we provide a physical interpretation to (∆*x*)^2^ in terms of the squared distance from the origin per generation. Let the distance 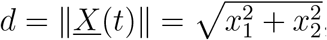, then

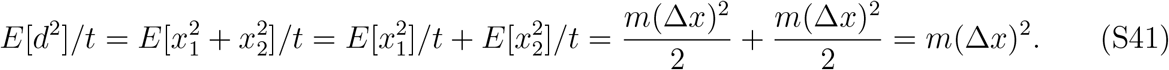

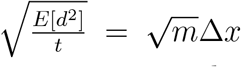 can be interpreted as the distance traveled by an individual after one generation, and sometimes is called the “dispersal” distance or the “root mean square distance”.

### 5.3 Diversity rates versus coalescent rates

For computational efficiency, the EEMS software uses a combination of the resistance distance model and within-deme “diversity rates” to approximate expected pairwise coalescent times, in which,

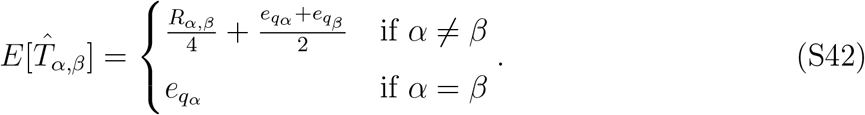

where *E*[*T̂*_αβ_] is the resistance distance approximation to the expected coalescent time between deme *α* and deme *β*, *e_q_α__* is the “diversity rate” in deme *α*, and *R_α,β_* is the resistance distance between demes *α, β* (Petkova et al., 2016). The diversity rates have no simple expression in terms of population-genetic parameters under the multi-deme coalescent model. As an alternative, diversity rates can be interpreted as reflecting average within deme heterozygosity since *e_q_* = *E*[*T̂*_*w*_] ∝ *H_α_* where the heterozygosity for deme *α* (*H_α_*) is defined as,

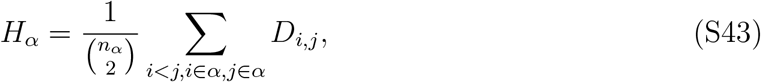

where *D_i,j_* is the average number of differences between (haploid) individuals *i* and *j*.

#### Migration and population sizes are identifiable in MAPS

MAPS models the recombination process using rates estimated from a recombination rate map. In this model, population sizes and migration rates can be inferred separately rather than as a joint parameter. Intuitively, the recombination rate serves an independent clock to calibrate estimates.

More formally, a statement of identifiability is a statement regarding the likelihood. MAPS models the expected number of lPSC segments shared between pairs of (haploid) individuals, and can be computed with an integral (14). The integral can be broken up into a product of two functions: a function describing the decay of PSC segments as a function of time (“recombination rate clock”), and the coalescent time probability density *f_T_i,j* (*t*) (15). The migration rates and population sizes only appear in *f_T_i,j* (*t*), and cannot be cannot be factored into parameters involving combinations of the migration rates and population sizes.

### 5.4 The prior

The structure of the prior closely resembles the prior in the EEMS method Petkova et al. (2016). The tessellation for the migration rates (*T_m_*) is encoded by a list (*<u>l^m^</u>, <u>m</u>, c_m_, μ_m_*) where *<u>l^m^</u>* are the locations of each cell, *<u>m</u>* the rates of each cell, and are vectors of length *c_m_* (i.e. number of Voronoi cells), and *μ_m_* is the overall mean migration rate. The Voronoi tessellation for the coalescent rates is *T_q_* = (*<u>l^q^</u>, <u>q</u>, c_q_, μ_q_*).

The location of each (unordered) Voronoi cell is distributed uniformly across the habitat,

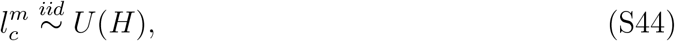

and the number of cells (a-priori) are drawn from a negative binomial distribution,

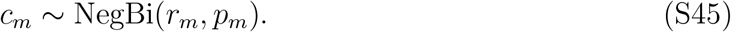

The effects of each Voronoi cell is normally distributed with variance *ω*^2^.

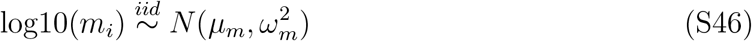

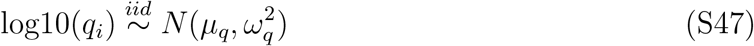

The probability of a particular (unordered) cell configuration is,

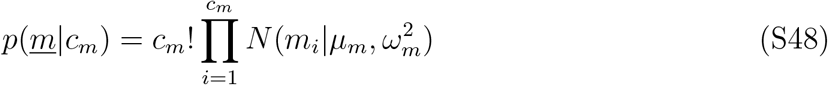

We assume,

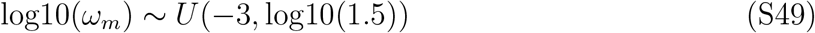

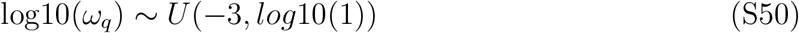

We set log10(2) as the upper bound for log10(*ω_m_*) so the *m* so the probability that it is within 3 orders of magnitude from the mean is 0.95 a priori, and we set log10(1) as the upper bound for log10(*ω_q_*) to restrict the population sizes so to be within 2 orders of magnitude from the mean with probability 0.95 a priori.

We assume,

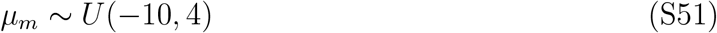

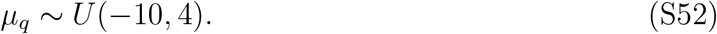

We place a uniform prior on the log of the mean rates to reflect that we are uncertain about the order of magnitude. Here, the data is highly informative of the mean, as a result, we can allow the support of the prior to vary by many orders of magnitude.

### 5.5 MCMC

#### Re-parameterization

We re-parameterize the model to improve mixing of the MCMC. We decouple the migration (or coalescent) rates from the mean rate (*μ*), and variance (*ω*) by introducing a new variable *e_i_*,

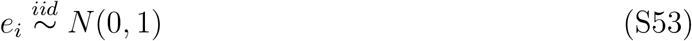

and the cell specific migration (or coalescent) rates are computed as,

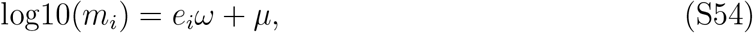

which allows us to update the magnitude of the parameters (*μ*) and the variance scale (*ω*) separately.

We add MH joint random-walk updates to *μ* and *e_i_* to ensure that 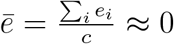. To do this, we jointly update *μ* and *e_i_* by,

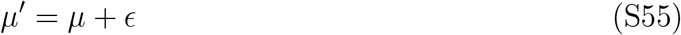

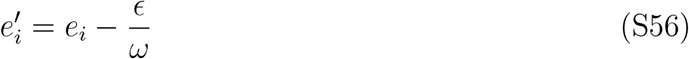

where *∊ ~ N* (0, 1). We do this for both the migration rates and population sizes.

#### Updating the number of cells

The number of cells change the dimension of the likelihood, and a result, we must use a Reversible Jump MCMC step so that the ratio of densities in the Metropolis-Hastings acceptance ratio is well-defined (Green, 1995). We choose to update the number of cells with a birth-death update (Stephens, 2000). Fortunately, in such a case, the updates reduce to standard Metropolis-Hastings because the dimension matching constant (i.e. the “Jacobian”) equals one (Petkova et al., 2016; Stephens, 2000). See equations S31 and S32 in Petkova et al. (2016) for formulas regarding the birth-death update. Here, we use nearly identical updates (with a slight modification).

When increasing the number of cells from *c* to *c* + 1 (i.e. a birth-update), we randomly choose a location uniformly across the habitat, and the new migration is proposed from a standard normal because our cell effects are standardized. In contrast, EEMS proposes cell effects migration to be normally distributed around a cell effect at a randomly chosen point in the habitat. Here we set, p(birth) = p(death) = 0.5 if the number cells *≥* 1, otherwise p(birth) = 1.

The acceptance ratio for a birth update (going from *c* cells to *c* + 1 cells) is

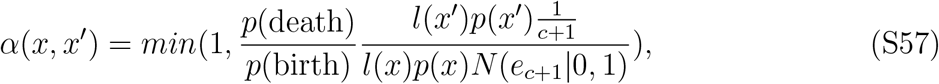

where *x* denotes the current state of the MCMC, *x′* the proposed state, *e*_*c*+1_ is the proposed cell effect drawn from a standard normal. Conversely, in a death-update, we randomly choose one cell uniformly to kill. In this case, the acceptance ratio for a death proposal (going from *c* + 1 cells to *c* cells) is

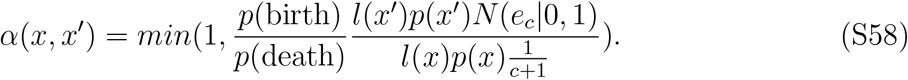

## 6 Supplementary Figures

**Figure S1:**
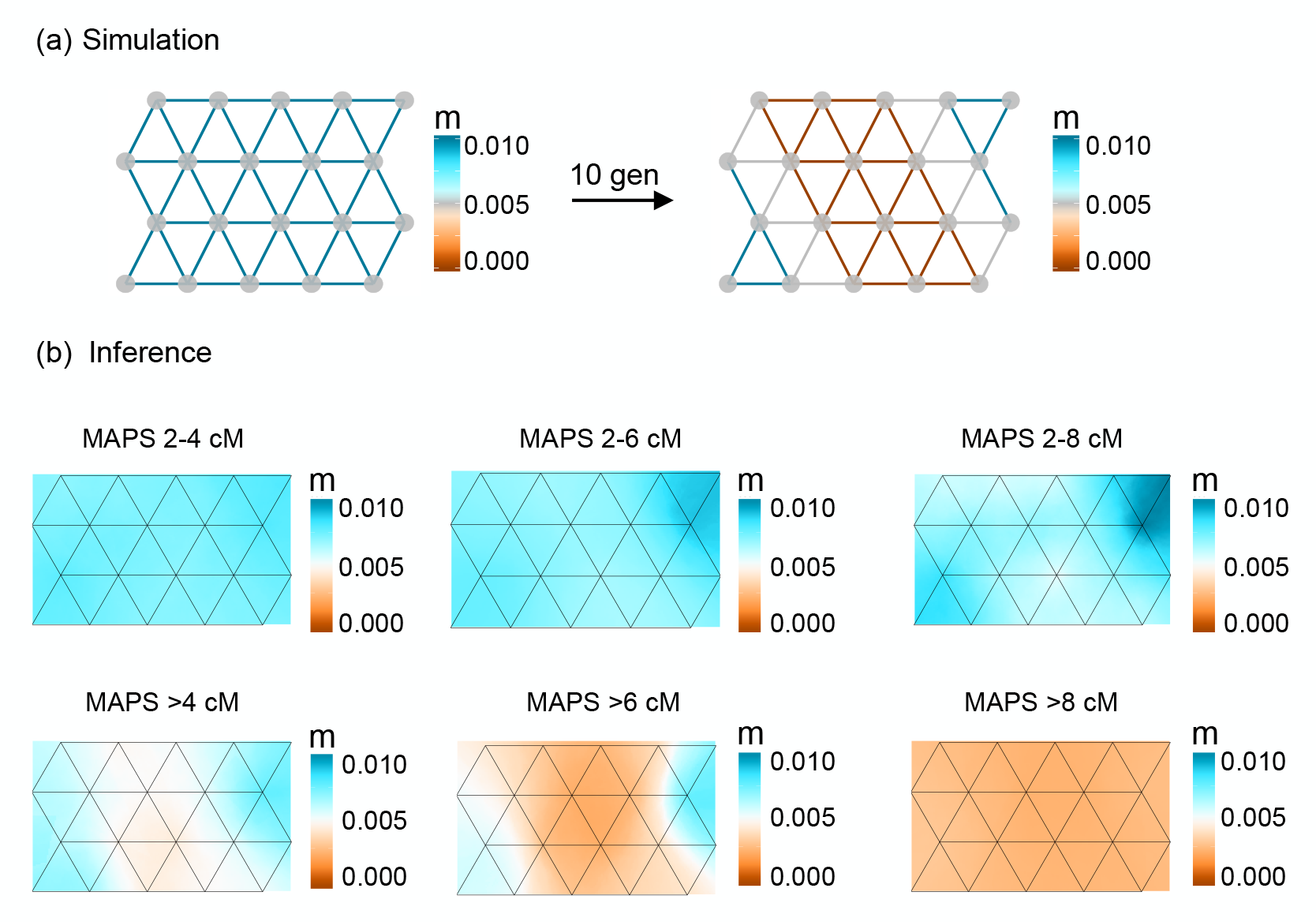
The performance of MAPS on a recent barrier scenario under different PSC length bins. Here, we investigate the ability of MAPS to detect a recent barrier (*<* 10 generations) for various PSC length bins (a) Simulation scenario. Population sizes were set to 10,000 per deme and 10 diploids were sampled per deme, replicating the conditions in Figure 2b. (b) Results for different PSC length bins. Length bins that encompass shorter segments (2-4cM 2-6cM 2-8cM) recover the higher uniform migration surface; while length bins with longer segments (>4, >6, >8) recover the recent ancestral barrier. For the last length scale (> 8cM), the signature of low migration extends across the habitat. The variation in migration rates is missed presumably because of the small number of shared segments at this length scale.

**Figure S2:**
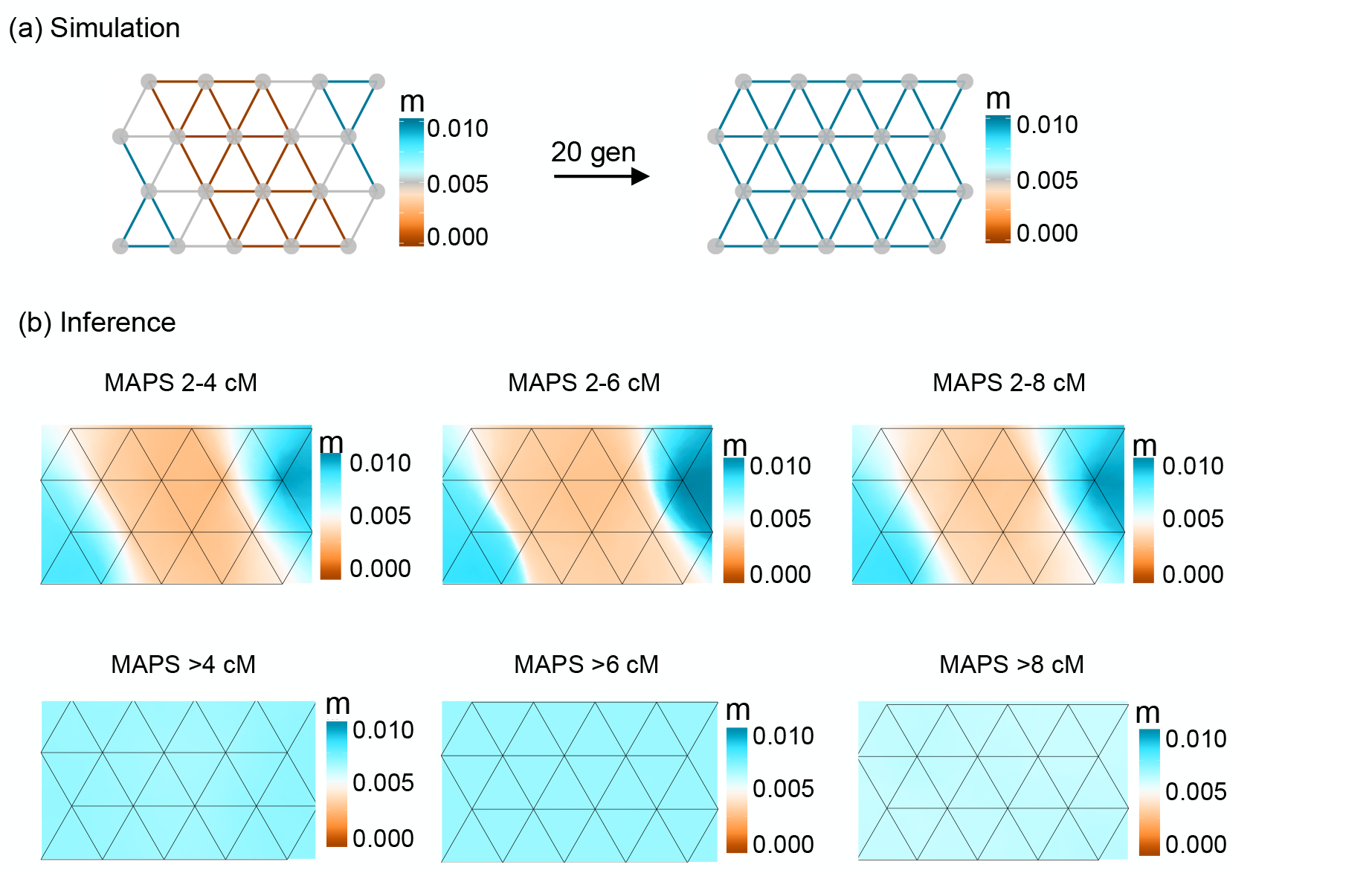
The performance of MAPS on a past barrier scenario under different PSC length bins. a) Simulation scenario. Population sizes were set to 10000 per deme and 10 diploids were sampled per deme, replicating the conditions in Figure 2c. (b) Results for different PSC length bins. Length bins that encompass shorter segments (2-4cM, 2-6cM, 2-8cM) recover the ancestral barrier; while length bins with longer segments (>4, >6, >8) recover the recent constant migration surface.

**Figure S3:**
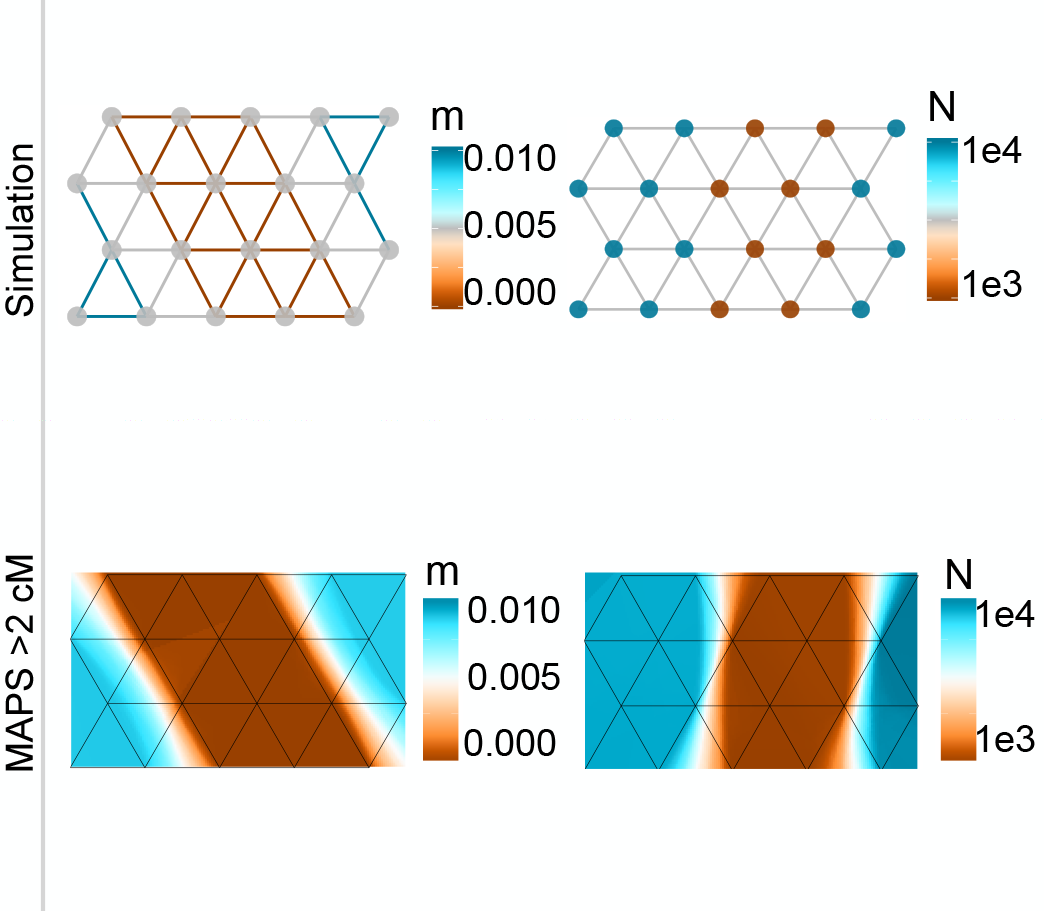
The performance of MAPS under a jointly heterogeneous migration rate and population size surface. a) Simulation Scenario. Heterogeneous population-sizes and migration rates (as shown) were simulated, and 10 diploid individuals were sampled per deme. (b) Results for PSC segments greater than 2cM are shown.

**Figure S4:**
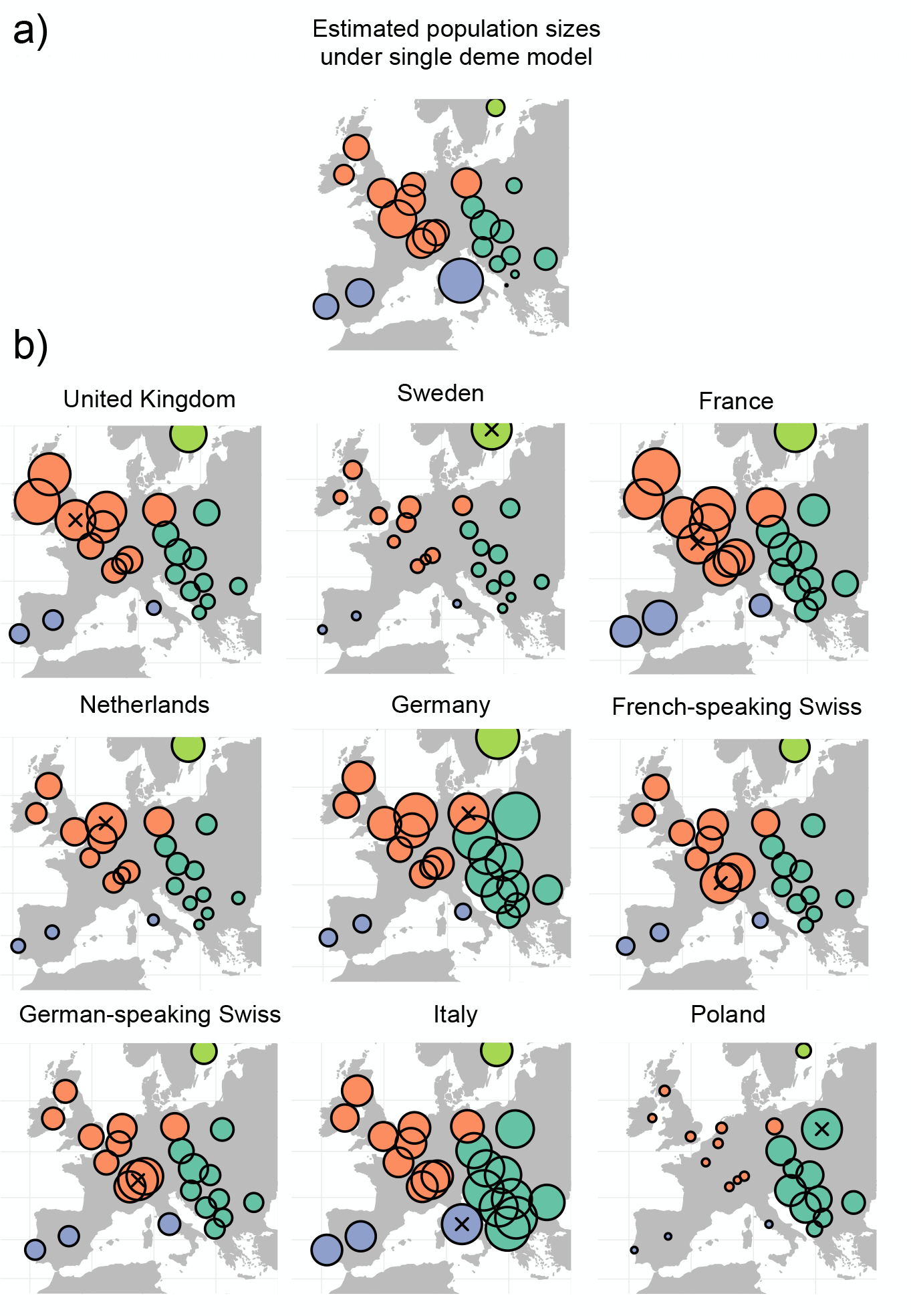
Visualizing normalized sharing of PSC segments that are 1-5cM. The color scheme is the same as used in Ralph and Coop (2013) where the colors give categories based on the regional groupings: W Western Europe, S Southern Europe, and E Eastern Europe (a) The average sharing within each sample locale is transformed to population sizes using the simple single deme estimator by Palamara et al. (2012). This transformation can be roughly summarized as to say that 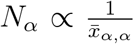 where *N_α_* is the effective population size in deme *α* and *x*_*α,α*_ is the average pairwise PSC sharing between individuals in deme *α*. (b) Similar to Ralph and Coop (2013), for each focal population (marked with an x), we plot the normalized average pairwise sharing between that population and all others (normalized by the average sharing within the focal population), i.e. if *α* is the focal population, we show 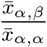 for each other country *β*.

**Figure S5:**
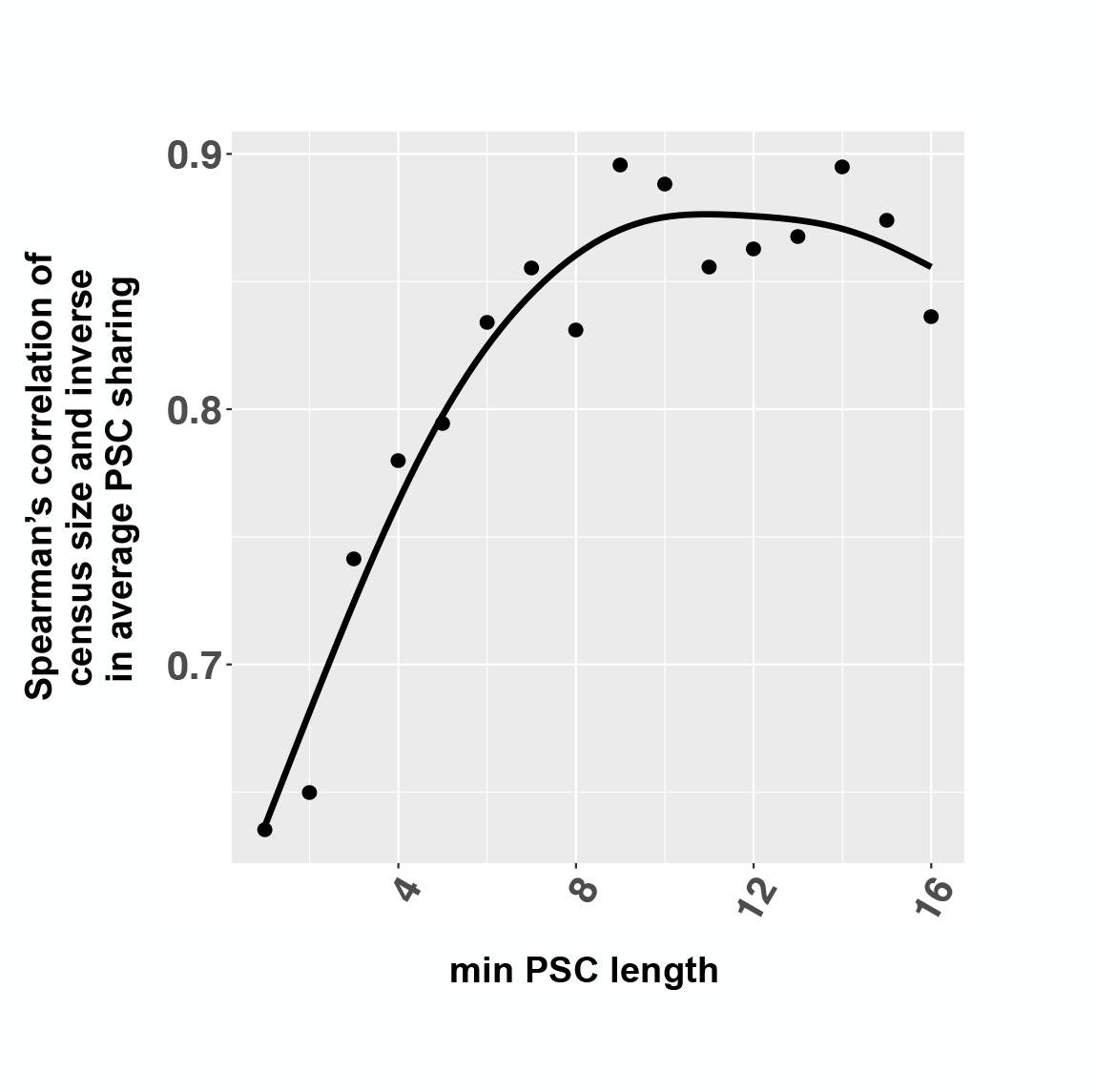
The correlation between census size and inverse average PSC sharing as a function of minimum PSC length considered. We use census size compiled from the The World Bank (2016) and National Records of Scotland (2011). The smooth black curve denotes the loess fit. Longer PSC segments correlate more strongly with census size than shorter PSC segments

**Figure S6:**
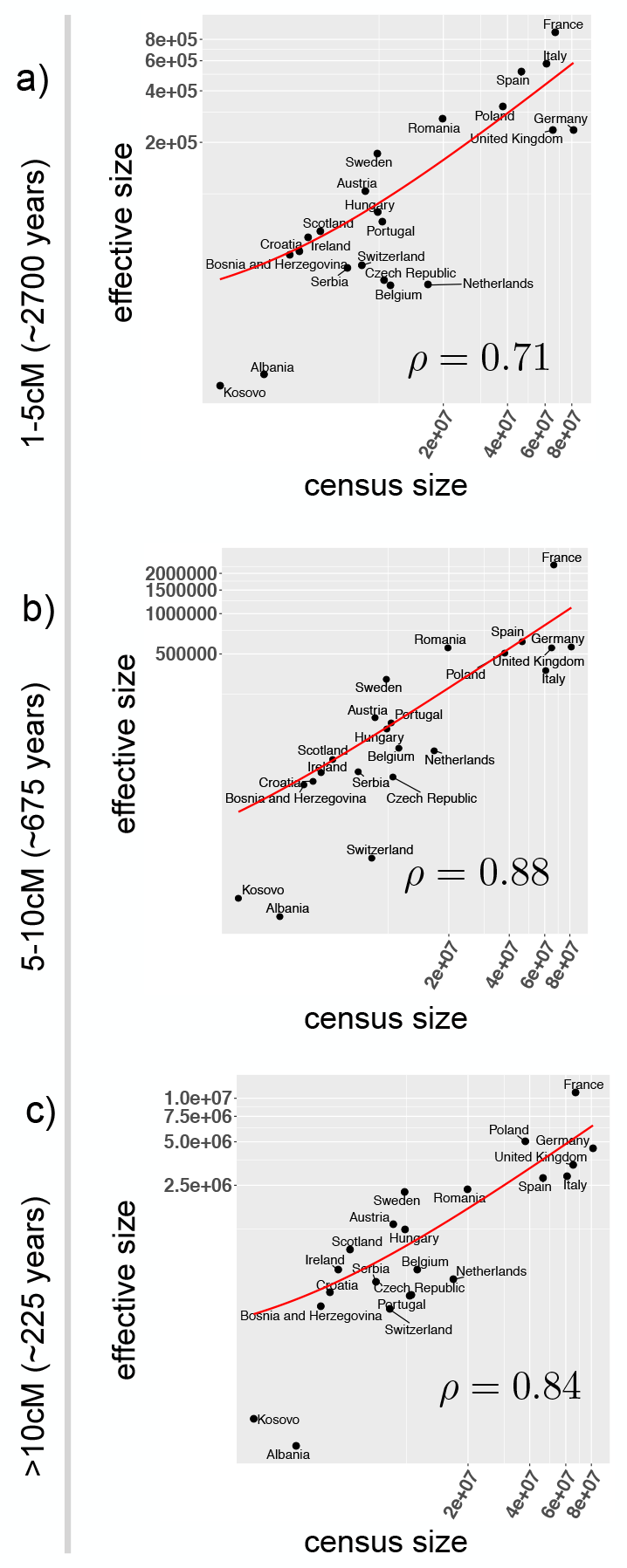
Census size versus MAPS estimated population sizes. Using the MAPS output, we estimate a total size per population by summing the estimated deme-level sizes across the area of each respective country (whether’s a deme’s location falls within a country was determined by querying The GeoNames Geographical Database). Finally, we plot the results on a log10 scale for different length scales (a) 1-5cM, (b) 5-10cM, and (c) >10cM. The red curve denotes the linear fit on the absolute scale. Note Kosovo and Albania as candidate outliers possibility because of cryptic relatedness artificially decreasing population sizes.

**Figure S7:**
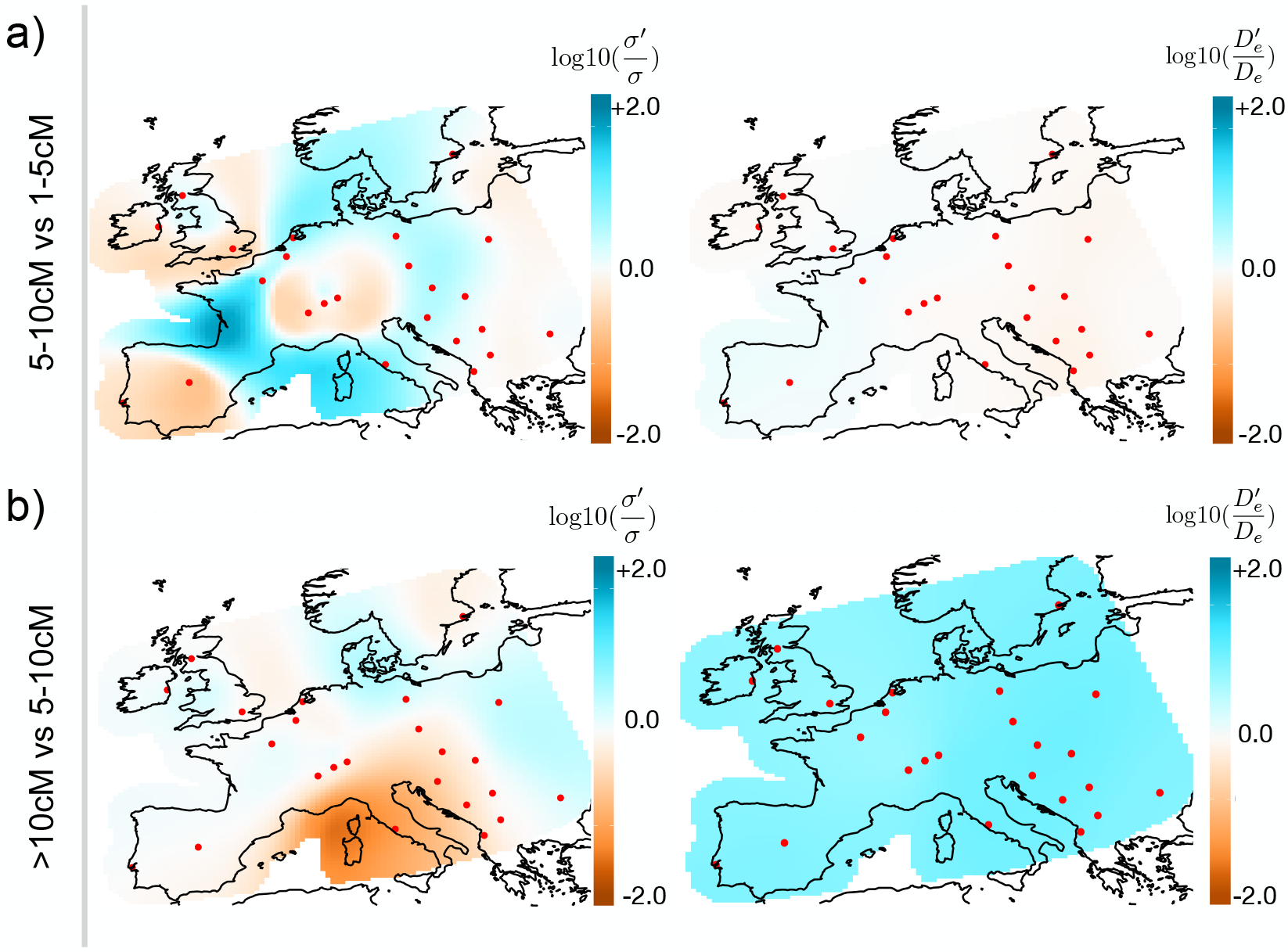
Plots of estimated average log10 differences in demographic parameters between adjacent time scales. (a) We plot estimates of 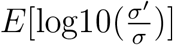 and 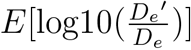 across the spatial habitat where *σ′*(*D′_e_*) denotes the dispersal rates (population densities) in the 5-10cM length bin and *σ* (*D_e_*) denotes the dispersal rates (population densities) in the 1-5cM length bin. (b) The results here are similarly plotted as above, however, the adjacent length scales are given by: 5-10cM and >10cM. The log10 differences are estimated in such a way so that the mean log10 difference is shrunk to zero. For example, for estimating dispersal in 5-10cM, we assume log10(*σ′*) = *E*[log10(*σ*)] + *∊* where *E*[log10(*σ*)] is estimated using PSC segments 1-5cM and *∊ ~ N* (0, *ω*^2^) is estimated from PSC segments 5-10cM. Consequently, the log ratio between dispersal rates from the two lengths bins is constructed to have mean zero *apriori* (i.e. *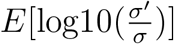* = 0).

**Figure S8:**
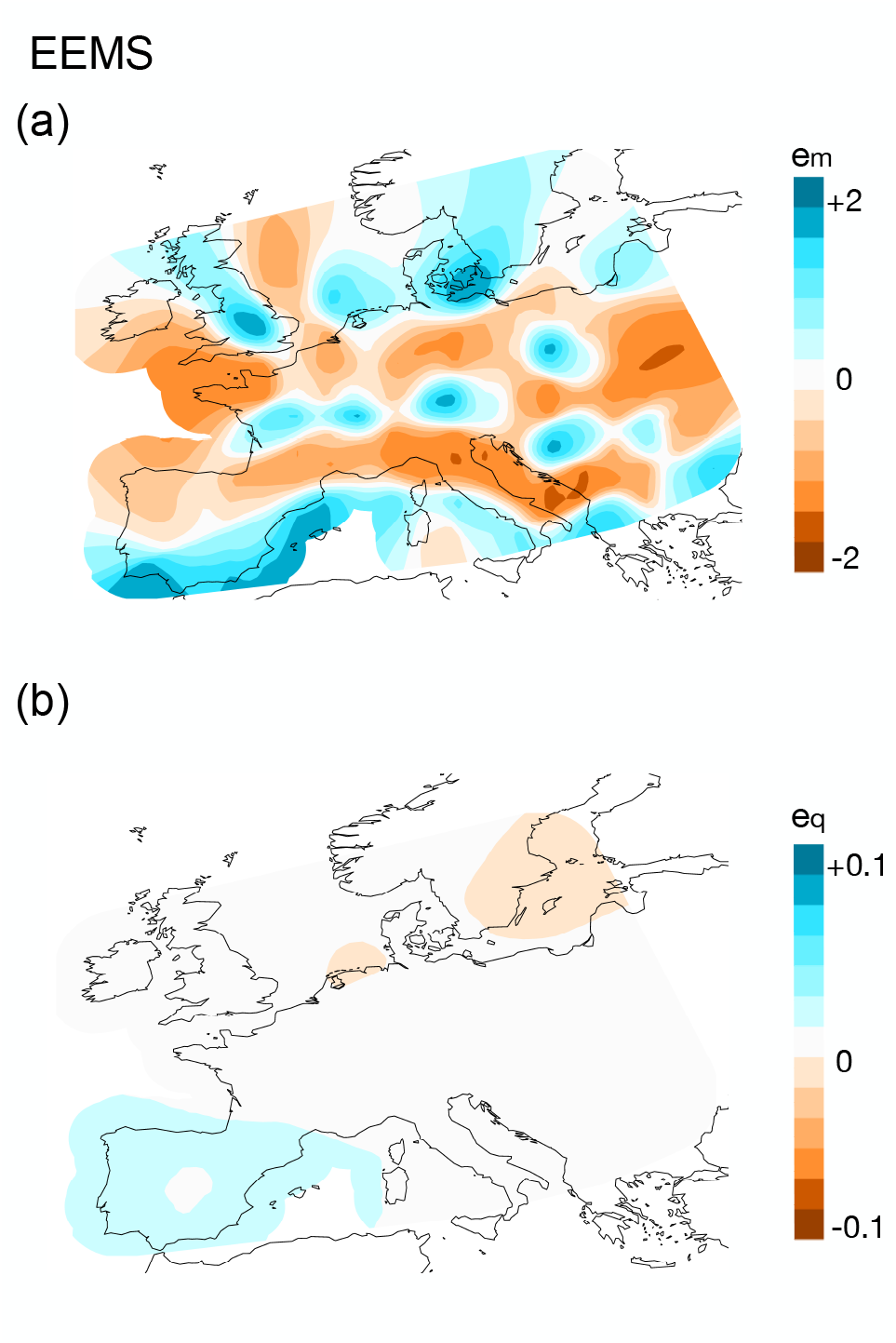
EEMS applied to the POPRES dataset. We apply EEMS to the same set of individuals as used in Figure 4 (see Methods). (a) The effective migration rates (b) The effective diversity rates. Here, we ran EEMS with 200 demes (as in Figure 4) with default parameters and averaged over 10 independent replicate chains. Each chain ran with 50e6 MCMC iterations, 25e6 set as burn-in, and we thinned every 5000 iterations.

**Figure S9:**
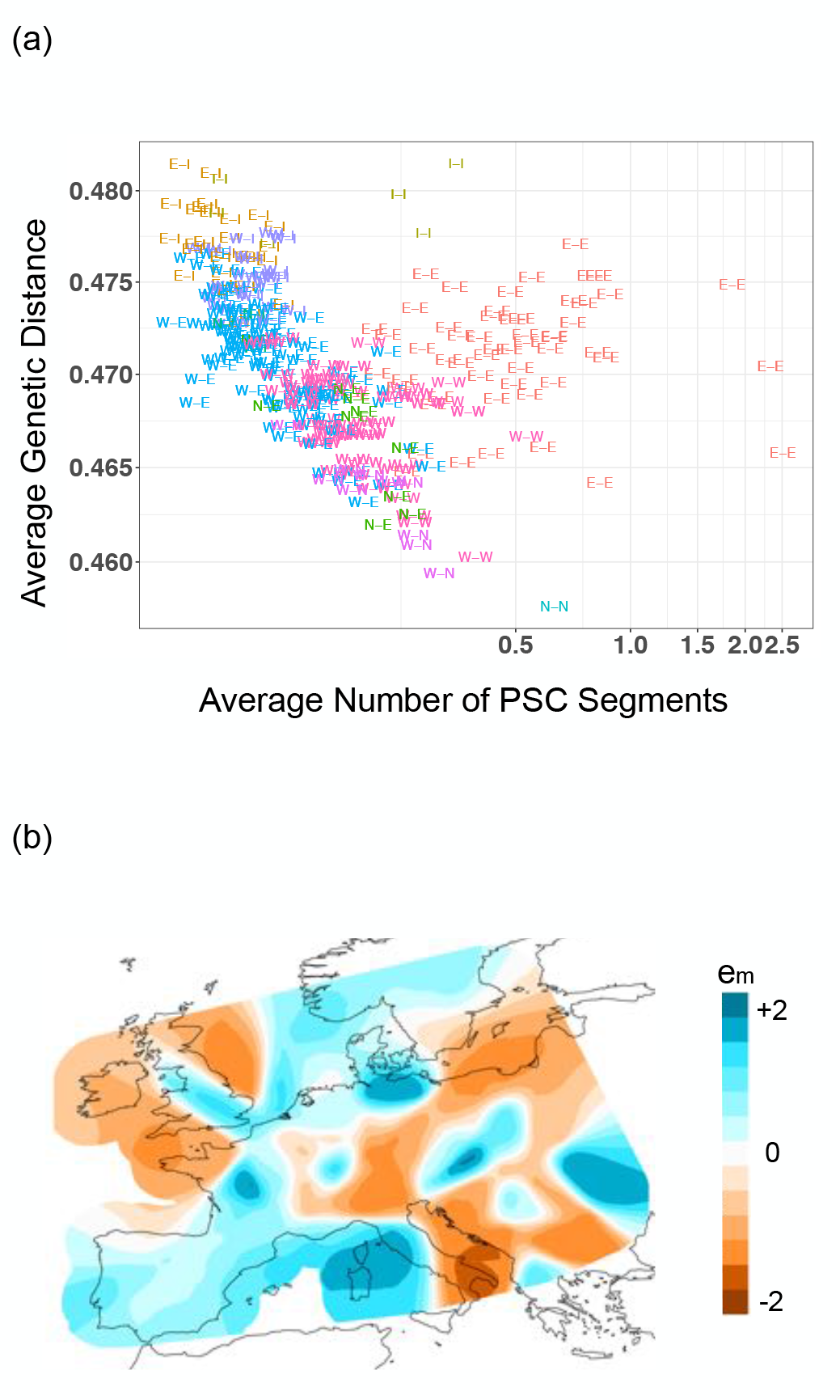
Genetic distance vs PSC sharing. (a) The averaged genetic distance (as used in EEMS) is plotted against the average number of PSC segments (> 1cM) for each pair of populations. Each point denotes a pair, the symbols represent groupings from Ralph and Coop (2013) (W Western Europe, S Southern Europe, and E Eastern Europe), and the colors represent the pair of regions. We see a negative correlation between the two summary statistics (Pearson’s *ρ* = −0.38, p-value = 7e-11), with the largest deviations occurring in comparisons between Eastern European populations. (b) EEMS results on PSC data transformed to a distance matrix. First, we encoded the PSC sharing statistics into a similarity matrix *S* such that *S_i,j_* is the number of shared PSC segments between samples i and j and *S_i,i_* is the maximum number of shared segments in the dataset (which we denote as *c*) to ensure *S* is a similarity matrix. Next, we transformed *S* to a genetic distance matrix *D* such that *D* = *c*11*^T^ − S* + *E* where *E ≈* 0 is a random genetic distance matrix of normal vectors with mean 0 and standard deviation of 0.01 added to ensure *D* is full rank. Finally, we applied EEMS to the distance matrix *D*. Though this procedure is heuristic, we see shared features between this surface and the MAPS dispersal surface shown in Figure 4.

